# The XyloPhone: democratizing access to high-quality macroscopic imaging for wood and other substrates

**DOI:** 10.1101/2020.08.02.233114

**Authors:** Alex C. Wiedenhoeft

**Affiliations:** Center for Wood Anatomy Research, Forest Products Laboratory, Madison, WI 53726, USA; Department of Botany, University of Wisconsin, Madison, WI 53706, USA; Department of Forestry and Natural Resources, Purdue University, West Lafayette, IN 47907, USA; Ciências Biológicas (Botânica), Universidade Estadual Paulista – Botucatu, São Paulo, Brasil; Department of Sustainable Bioproducts, Mississippi State University, Starkville, MS, USA

**Author notes:** primary affiliation.

## Abstract

One rate-limiting factor in the fight against illegal logging is the lack of powerful, affordable, scalable wood identification tools for field screening. Computer vision wood identification using smartphones fitted with customized imaging peripherals offer a potential solution but to date, such peripherals suffer from one or more weaknesses: low image quality, lack of lighting control, uncontrolled magnification, unknown distortion and spherical aberration, and/or no access to or publication of the system design. To address cost, optical concerns, and open access to designs and parameters, I present the XyloPhone, a 3D printed research quality macroscopic imaging attachment adaptable to any smartphone. It provides a fixed focal distance, exclusion of ambient light, selection of visible light or UV illumination, uses the lens from a commercially available loupe, is powered by a rechargeable external battery, is fully open-sourced, and at a price point of less than 110 USD is a highly affordable tool for the laboratory or the field, and can serve as the foundational hardware for a scalable field deployable computer vision wood identification system.

## Introduction

### The need for forensic wood identification to combat illegal logging

Calls for various forms of scientific timber testing to combat illegal logging and prevent fraud are well established in the literature (Johnson and Laestadius 2011; Dormontt et al. 2015; Expert Group UNODC 2016; Lowe et al. 2016; Schmitz et al. 2019; Wiedenhoeft et al. 2019; Schmitz et al. 2020). Answers to those calls necessarily focus on the underlying biological variation inherent in wood itself, and therefore address questions not of paperwork or permits, but rather of the botanical identification (“species”), the geographic origin, or individualization (log to stump, board to log, etc.) of wood. There are two broad sources of that variation in wood that are relevant for scientific timber testing, molecular variation, and structural or anatomical variation.

Methods interrogating molecular variation in wood are almost all limited to a laboratory setting and for the foreseeable future cannot be expected to have practical field deployability (e.g. mass spectrometric methods, DNA barcoding, DNA-based individualization, DNA-based population assignment, other chemometric methods such as LIBS, pyrolysis mass spectrometry, DART-ToF), but what these methods lack in field relevance, they may provisionally make up for in the promise to resolve: species-level identification, identification of geographic origin, and individualization. The one molecular-based technique with a demonstrated potential for field deployment is near infrared spectroscopy (Snel et al. 2018). Limits to field deployability of NIRS are primarily the comparatively narrow range of taxa for which published models exist (Pastore et al. 2011; Bergo et al. 2016; Soares et al. 2017; Silva et al. 2018), and the rather high per-unit costs for field equipment (~ at least 10k USD).

Methods making use of the structural variation in wood include human-based wood identification using various forms of macroscopy and microscopy in the laboratory (Koch et al. 2011; Koch et al. 2018; Gasson et al. 2011; He et al. 2020), human-based field identification of wood using a loupe (Miller et al. 2002; Miller et al. 2004; Miller et al. 2005; Wiedenhoeft 2011; Ruffinatto et al. 2015; Yin et al. 2016; Arévalo et al. 2020; Ruffinatto and Crivellaro 2020), and computer vision wood identification which can be implemented in the laboratory or in the field (Khalid et al. 2008, Martins et al. 2013; Filho et al. 2014; Figueroa-Mata et al. 2018; Ravindran et al 2018; Ravindran et al. 2019; de Andrade et al. 2020; Lopes et al. 2020; Olschofsky and Kohl 2020; Ravindran and Wiedenhoeft 2020; Souza et al. 2020). Wood anatomy, even with full access to reference collections and microscopic modes of evaluation, is rarely accurate to the species level when performed by human analysts (Gasson 2011), and field level wood identification is typically expected to be less accurate still. Research using machine learning in wood identification demonstrates that species-level resolution may be possible based on wood anatomy, either by employing machine learning in conjunction with human expertise (Esteban et al. 2009; Esteban et al. 2017; He et al. 2020) or by computer vision wood identification systems operating on images alone, mostly restricted to the laboratory (Martins et al. 2013; Filho et al. 2014; Rosa et al. 2017; Figueroa-Mata et al. 2018; Ravindran et al 2018; Souza et al. 2020).

### Field-deployable computer vision wood identification (CVWID) systems

Only three potentially field-deployable computer vision wood identification (CVWID) systems have been published to date (de Andrade et al. 2020; Lopes et al. 2020; Ravindran et al. 2020), with a fourth available commercially but lacking peer reviewed literature explicitly subtending it (https://www.xylorix.com). Three of the four systems use images collected from smartphones with image quality varying from quite poor, showing obvious and extreme spherical aberration (https://www.xylorix.com/wood-directory/), to good (de Andrade et al. 2020), or with image quality essentially impossible to evaluate (e.g. Figure 1, Lopes et al. 2020) based on the failure to adequately prepare specimen surfaces for imaging wood anatomical features. The fourth CVWID system, the XyloTron, requires custom imaging hardware and a laptop (Ravindran et al. 2020) to conduct real-world, in-field, on-device inference (Ravindran et al. 2019) and is the only system to report actual field use and inference in real-time.

**Figure 1.**
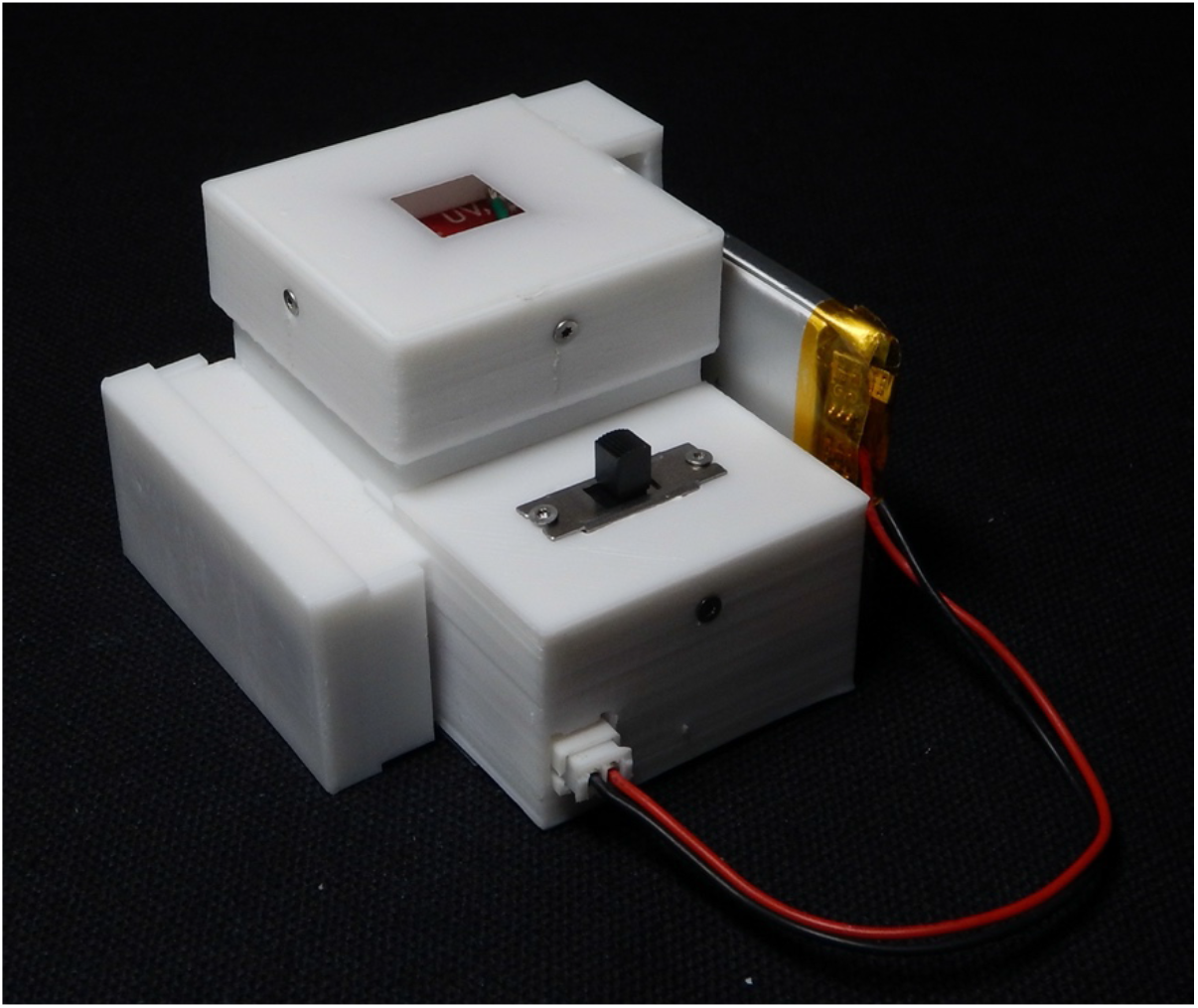
The XyloPhone printed in white polymer to facilitate photography. When mated to a phone-specific mounting plate, the XyloPhone can be used with any smartphone. As noted in the Supplementary Information, printing the XyloPhone in black polymer provides the best performance.

### Imaging hardware design considerations for field-deployable CVWID systems to combat global illegal logging

In addition to topics covered in the conceptual overview of imaging wood for CVWID (Hermanson and Wiedenhoeft 2011), there are three broad factors that inform design and implementation imaging hardware for field-deployable CVWID systems: a form factor related to necessary operator expertise; cost of the system; and, optical and imaging parameters.

#### Necessary operator expertise and device form factor

For all the CVWID systems in existence to date, the operator must cut or polish the transverse surface of a wood specimen such that the wood anatomical features are exposed. Failure to do this well (e.g. Figure 1 in Lopes et al. 2020) could result in spurious results when training a model – assuming that a system is purporting to identify the woods based on wood anatomy, if the characteristic wood anatomy is not captured in the images no amount of machine learning can generalize a model, nor can a model built from good data properly identify an image during the inference phase from a specimen prepared in such a way that does not show the anatomy. Beyond this manual skill (easily taught to undergraduates in wood identification classes around the world and to law enforcement officers when training for field wood identification), operator expertise mostly involves orienting the imaging hardware on or near the surface of the wood and capturing an in-focus image. Imaging hardware should therefore be easy to position on the substrate, should be robust to small bumps or perturbations, should easily establish a focal distance, should be small enough to be convenient without being too small and fragile, and for phone-based systems should be easy to attach reliably to the phone in the same position each time.

#### Cost of the system

The XyloTron platform (Ravindran et al. 2020) is able to image and identify both wood and charcoal, is field-portable, and costs ~1300 USD to build the XyloScope, and then another several hundred USD for a suitable laptop for storage, power, visualization, and inference, but remains the *de facto* gold standard for standardized macroscopic wood imaging. The per-unit cost for a XyloTron is approximately 1800 USD, which while quite scalable compared to training humans or laboratory-based approaches, but remains impractical for many global contexts where its functionality could be valuable. The imaging systems described in Verly Lopes et al. (2020, the Ollo Clip, ~60 USD), de Andrade et al. (2020, unspecified lens costing ~3 USD), and the Xylorix system (29 USD) are clearly much more affordable and therefore vastly more scalable. With these latter systems, a bring-your-own-device model is implied, leveraging the comparative ubiquity of smartphones.

#### Optical and imaging parameters

Whether concerned with field imaging for real-time identification or collection of reference images, optical and image quality metrics should be important factors guiding the design of any new system, but to date reports on new systems have not published such details. Ravindran et al. (2020) specify the exact camera and lens in the XyloTron and the fixed field of view of each image, and de Andrade et al. (2020) specify the camera zoom, image size in pixels, and total image field of view in millimeters of their images, whereas Lopes et al. (2020) and Xylorix provide no such detail. Given the ubiquity of zoom functionality in smartphone camera apps, reporting the actual size of the resultant field of view is a critical factor. It would also be valuable to report the optical resolution of a system, the geometric distortion across a reference image, and the spherical aberration of the system. Such data facilitate comparisons between systems, as well as possibly providing useful information for post-processing images to correct systematic errors and to create effective data augmentation strategies for training identification models.

Also important is controlled illumination of the substrate. Of the works cited, only Lopes et al. (2020) make no effort to control illumination of the wood surface. The XyloTron (Ravindran et al. 2020) and the work by de Andrade et al. (2020) both exclude ambient light and provide controlled illumination, and the Xylorix system has a rechargeable LED lighting array to provide light, but the translucent hood between the lens and the specimen could permit the entrance of ambient light into the system. Additionally, the XyloTron is the only system to allow a user to select either visible light or UV illumination of the substrate. UV illumination permits the detection of wood surface fluorescence and can be useful for discriminating between woods with similar wood anatomy under visible light.

These factors are important not only as basic system parameters, but because depending on the image-based machine learning strategies employed to develop classification models, if distortion is present in an image, features of a given size in the center of the image will appear larger or smaller in the corners. With spherical aberration of sufficient severity, features with fine spatial scale will cease to be observable at the margins or corners of the images. Systematic error of these types endemic to a foundational data set could limit the accuracy or robustness of CVWID models developed from such data, depending on how such models are constructed.

For example, prior work in my laboratory (Ravindran et al. 2018; Ravindran et al. 2019; Ravindran et al. 2020; Ravindran and Wiedenhoeft 2020) uses multiple image patches from across the parent image to train classification models. If hardware-induced systematic error is present across the images at a scale or severity equal to or greater than the magnitude of image augmentation measures taken to control for such error, the resulting models could be less accurate, less robust, or even spurious. The garbage-in, garbage out (GIGO) rule of computer science can be one step more insidious in CVWID – garbage in, fiction out (GIFO), for example, CVWID models that claim species level accuracy over 98% in large models with many taxa that are considered inseparable by light microscopy, or models where the anatomical features were not observable in the parent images. GIFO is dangerous because the fiction-out might seem believable, even when too good to be true. The only way to ensure that *in silico* models have any appreciable real-world relevance (whether CVWID or other modalities) is through field testing that verifies the results forensically. Though it should go without saying, perhaps it needs to be said – collecting high-quality images is better than collecting low-quality images. It is trivial to add noise, distort, blur, or otherwise post-process high-quality parent images, but the reverse is not true.

### Access to the system parameters

A final factor when considering imaging hardware for a CVWID system is the relative access that a user will have to the details of the system itself. Open source systems (such as the XyloTron) allow end users to adapt the hardware for their specific use-cases as needed, whereas closed or commercial systems do not offer such access. Lack of access in this way forces a user to accept all design constraints whether or not they serve well for a specific use-case, which may reduce or constrain the utility of the CVWID system.

To address above noted concerns, in this paper I present the XyloPhone (Figure 1), an open source, 3D-printed research quality macroscopic imaging attachment adaptable to any smartphone. It provides a fixed focal distance, exclusion of ambient light, selection of visible light or UV illumination, uses the lens from a commercially available loupe, is powered by a rechargeable external battery, and at a price point of less than 110USD is a 12-fold price reduction over the XyloTron, while delivering comparable image quality. To document the efficacy of the XyloPhone I present comparative data on distortion, maximum resolution, and spherical aberration, as well as example images taken with the XyloTron, the XyloPhone on two different smartphones (an iPhone and a Samsung Android phone), and the Ollo Clip system on an iPhone.

## Materials and Methods

Smartphones: Two phones were used to collect data on field distortion and spherical aberration, a Samsung Note 5 running Android version 7.0, and an iPhone XS Max, running iOS 13.4.1. The native Android camera app was used on the Samsung Note 5, but on the iPhone the ProCamera app from the App Store was used to collect images. Images were saved in the JPEG format with square aspect ratio, maximum resolution and minimum compression.

Hardware configurations: All calibration and reference images were collected by mounting the phone + optical and lighting array (either XyloPhone or Ollo Clip + XyloPhone lighting array) to a 3D-printed phone holder that was secured to an x/y/z micromanipulator mounted to a stereomicroscope stand (Figure 2). This readily permitted the fine adjustments necessary for collecting reference images of the various targets for measuring optical performance, but is not required for wood image data collection or field use. This system was also used to collect the images of wood specimens in Figure 4 in order to capture substantially the same locations in the reference blocks.

**Figure 2.**
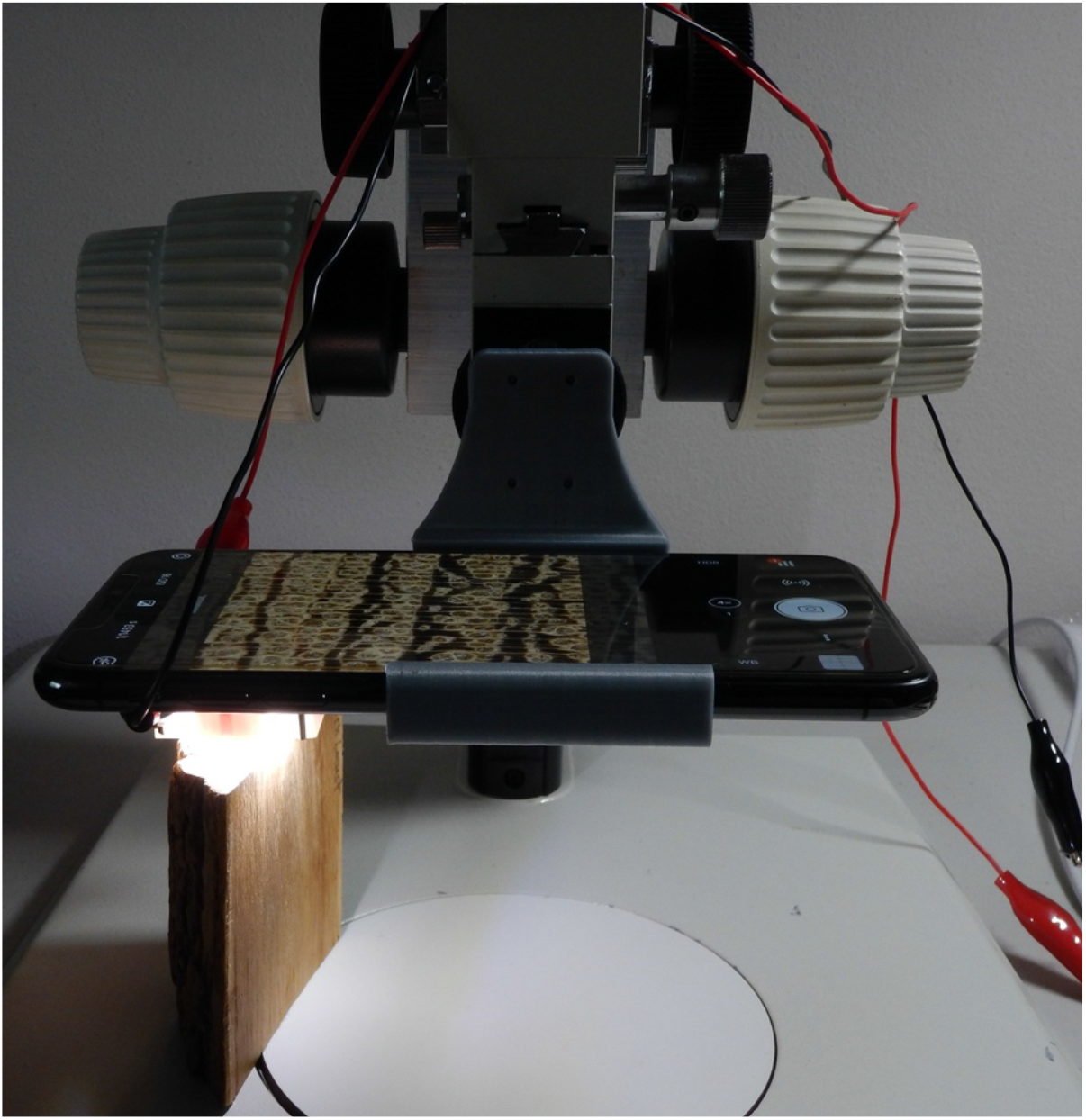
The 3D printed phone holder mounted to an x/y/z micromanipulator affixed to a stereoscope stand. The Ollo Clip + XyloPhone lighting array is shown illuminating a specimen of *Robinia pseudoacacia.*

Ollo Clip hardware: To collect comparable metrics for the 14X Ollo Clip lens I developed a 3D printed custom holder for the lens that mated with the iPhone and positioned the lens identically to the XyloPhone for scalar, field distortion, and spherical aberration measurements. Initial observations (data not shown) indicated that uncontrolled lighting with the OlloClip negatively affected the image quality. In order to maximize the validity of the comparisons between the XyloPhone and the OlloClip and to afford the OlloClip the maximum possible performance, I designed a jig (Figure 3) to position the XyloPhone lighting array in an equivalent position with the OlloClip, and this was used to collect all OlloClip images.

**Figure 3.**
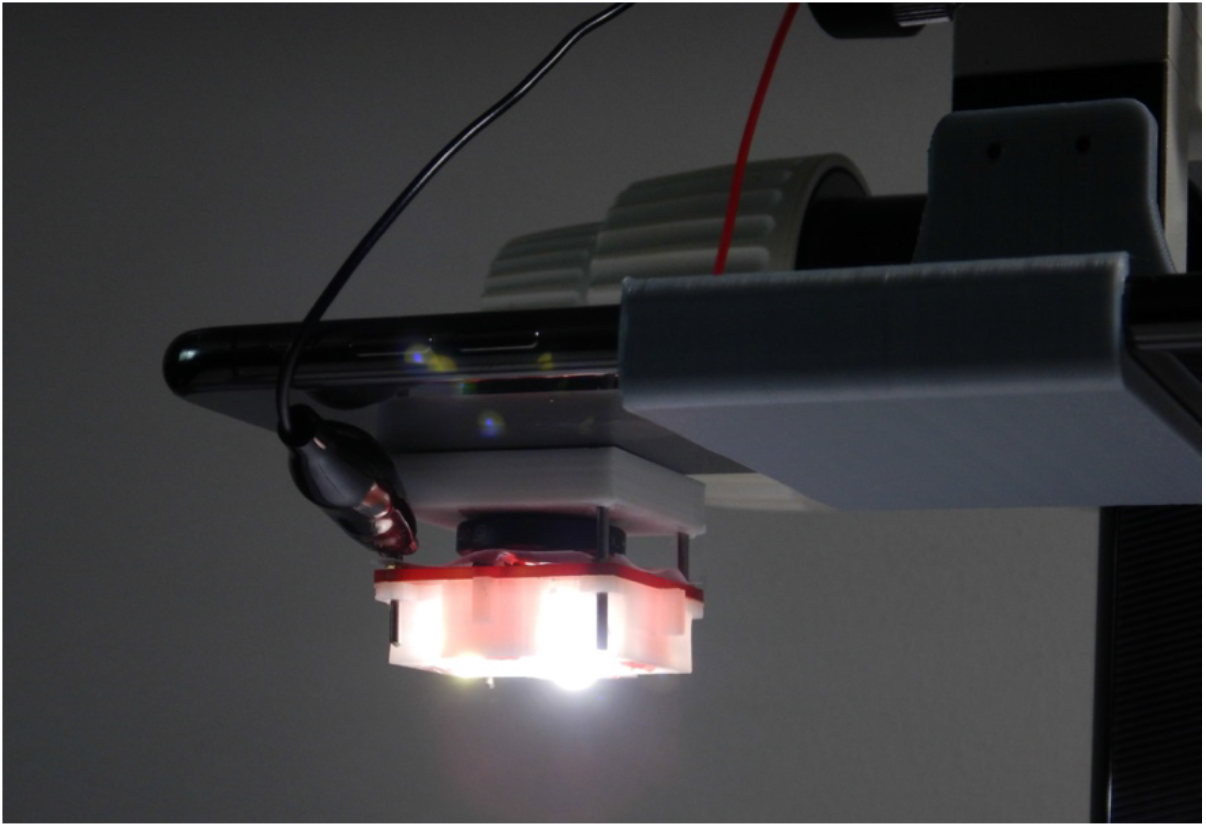
OlloClip lens jig and XyloPhone imaging array to collect OCi data.

**Figure 4.**
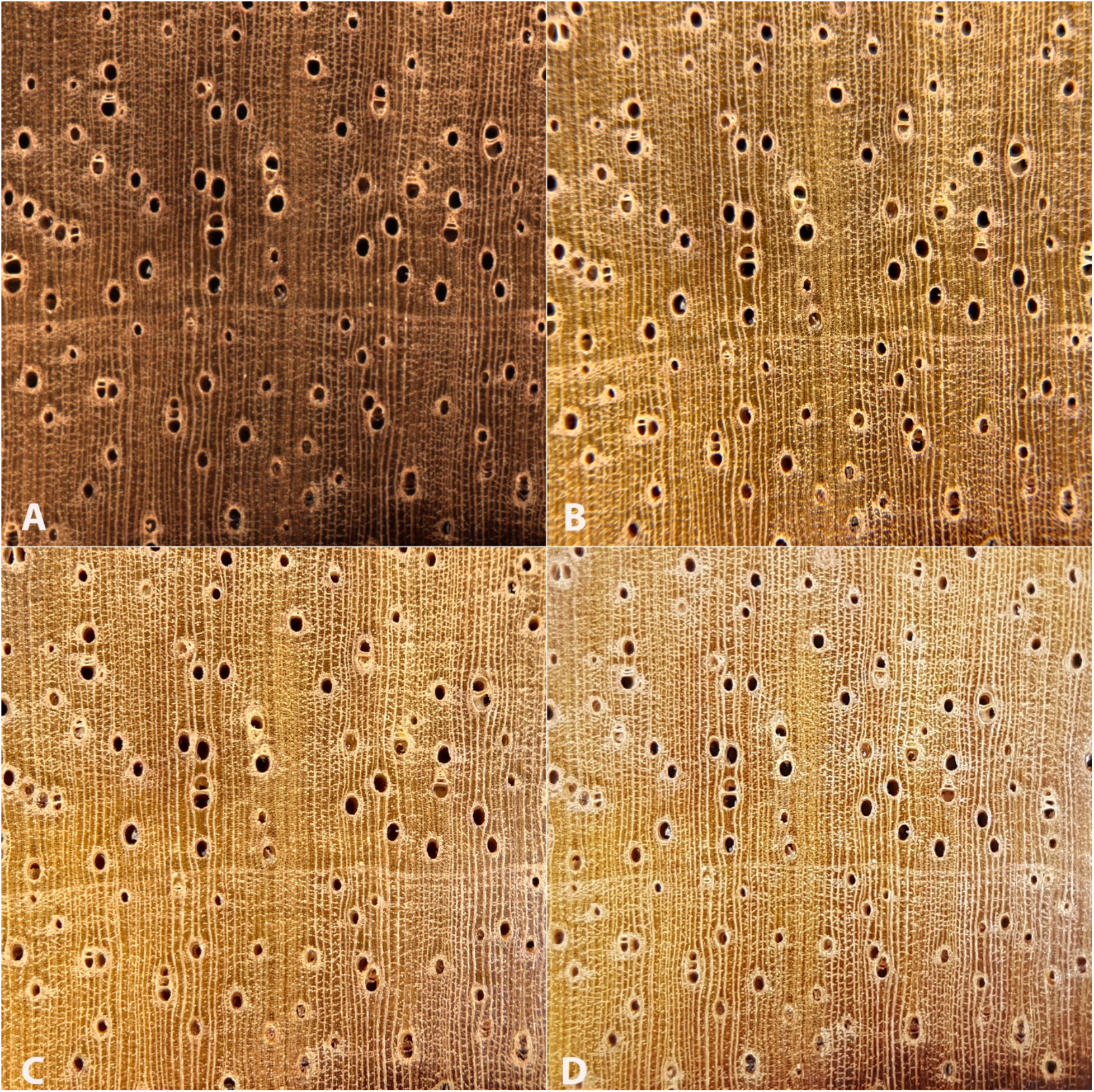
Macroscopic images of *Dalbergia retusa* sapwood of substantially similar fields of view. A, XT image. B, OCi image. C, XPi image. D, XPs image. Note the loss of focus (spherical aberration) in the corners of image B, and to a lesser degree D. The color balance of the XT image is set against a white balance card (Ravindran et al. 2020), whereas the white balance in the smartphone images was allowed to be controlled by the camera app.

Field of view: The size of the field of view was determined by capturing a reference image of the metric divisions of an Edmund Optics dual axis metric/English stage micrometer (#59277-0021) at each in-app camera zoom step to bracket the XyloTron’s field of view (~6.35mm × ~6.35mm, Ravindran et al. 2020). Resulting images were opened in Mac OS Preview and a bounding rectangle was selected from the leftmost edges of two reference values, and then nominal um per pixel were calculated from those values.

XyloTron as baseline: Because the XyloTron (Ravindran et al. 2020) is already used in a number of countries around the world (Ravindran et al. 2019; Arevalo et al. submitted, distribution of XyloTrons by Wiedenhoeft), we use the XyloTron image size as the baseline for comparison throughout the manuscript. For each device combination (smartphone + imaging/lighting hardware), the camera zoom level producing a field of view nearest to but not smaller than that of the XyloTron was chosen for data collection, resulting in an image that could be cropped and resized to be equivalent to the XyloTron field of view without need to alter the XyloTron image. The device combinations evaluated are the XyloTron (XT), the XyloPhone + iPhone (XPi), the XyloPhone + Samsung (XPs), and the Ollo Clip 14x + iPhone (OCi). These abbreviations are used throughout the balance of the manuscript.

Field distortion: Measurements were made on images of an Edmund Industrial Optics dot grid target (cert. 46250, s.n. 0000-0307). Images of the smallest dot grid (0.25mm diameter, 0.50mm spacing) were taken using the in-app camera zoom function to bracket fields of view as for determining field of view size. A XT image of the dot grid was also taken. Images were processed in ImageJ to determine the area per dot and total dot area of 3×3 patches of dots from the center of each image, and a 3×3 patch in each corner of each image, where distortion should be greatest.

Determination of dot diameter: Equation 1 was used to calculate the estimated diameter based on the average dot area determined in the distortion measurements and the μm per pixel resolution determined in the field of view measurements. The known dot diameter is 250 um

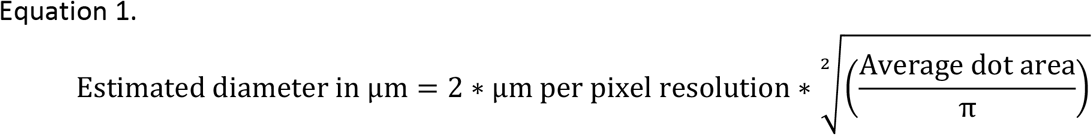

Image analysis: Images were processed in ImageJ v1.53c with the following steps: Adjust ≫ 8 bit; Adjust≫ Threshold (lower = 0, upper = 60); Analyze≫ Analyze particles; 5000-inifinity for size, Circularity 0.00-1.00. Measured properties included: Area; with limit to threshold checked. Ultimately, area was used as the metric from which calculate distortion, comparing the central 3×3 dots to 3×3 dots in each corner.

Determination of spherical aberration: An Edmund Optics NBS 1963A (#85276) positive target was used to determine the maximum resolvable line pairs per millimeter (lpmm) for each hardware configuration. Images of a range of lpmm were taken in the center of each field of view, and in each corner. Images were standardized to 100% zoom (in-silico) to evaluate the clear separation of line pairs, the maximum resolvable number of line pairs was recorded, then the mean value for each position (center, upper left, upper right, lower left, lower right) was calculated.

## Results

### The XyloPhone

The XyloPhone weighs ~93 grams (individual 3D printing parameters (e.g. infill density and pattern) and polymer choice will affect this), with another 10-20 grams contributed by the phone plate, for total of less 115 grams. With an approximate volume of 112 cm^3^ the XyloPhone is lightweight, small, and highly portable. The 1000mA rechargeable lithium-ion battery provides over 7.5 hours of continuous illumination and can be recharged with the in-device charger connected to any powered USB port on a computer or any DC 5V cell phone charger.

### Human-user evaluation of images

Figure 4 shows an image of substantially the same spot on a specimen of *Dalbergia retusa* sapwood. This taxon was chosen because its diffuse-in-aggregate parenchyma and numerous, narrow rays can fail to be resolved clearly when spherical aberration is prominent, as with the OCi (Figure 4, B) and the XPs (Figure 5, D). The XT image (Figure 4, A) appears to the naked eye to have consistent focus across the field of view, as does the XPi image (Figure 4, C).

**Figure 5.**
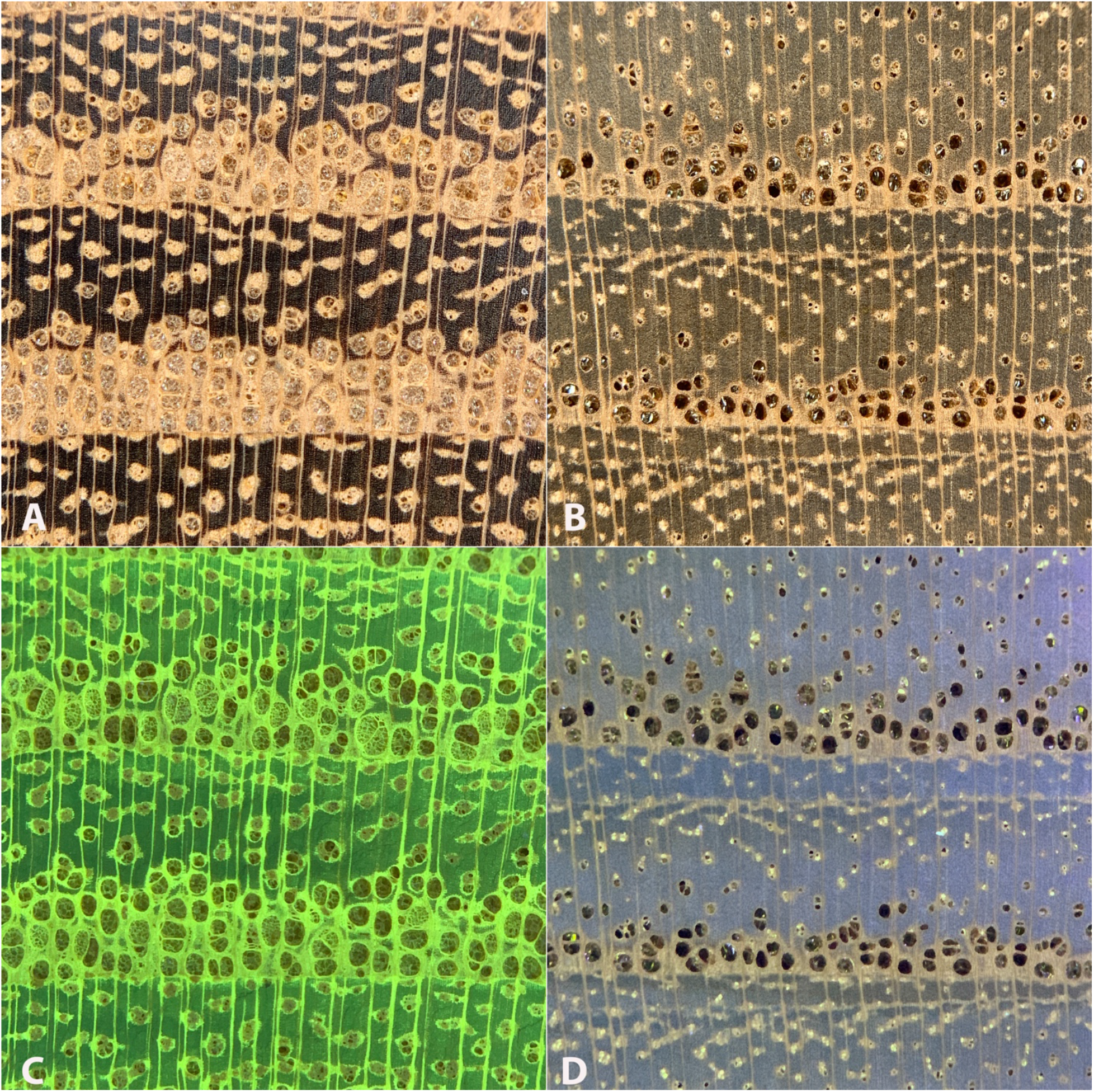
Free-hand XPi images of *Robinia pseudoacacia* (A, C) and *Morus rubra* (B, D) with visible light illumination (A,B) and UV illumination (C,D). Note the clear yellow-green fluorescence of *Robinia* in C. All images are 6,742 um on a side.

The visible light and UV light illumination options of the XyloPhone are demonstrated in Figure 5, which shows side-by-side images of *Robinia pseudoacacia* and *Morus rubra*. The heartwood fluorescence in *Robinia* is clearly visible.

### Quantitative evaluation of the optical properties of the XyloPhone, XyloTron, and Ollo Clip

Table 1 lists details for each of the hardware configurations studied here. Distortion ranged from 7% barrel distortion for the OCi, to 3% pincushion distortion for the XPs, to less than 1% pincushion for the XT and the XPi.

**Table 1.** Quantitative aspects of optical system performance for the Ollo Clip + iPhone (OCi), the XyloPhone + iPhone (XPi), XyloPhone + Samsung (XPs), and the XyloTron (XT). C = center of image, UL = upper left corner of image, UR = upper right corner of image, LL = lower left corner of image, LR = lower right corner of image.

|  | Image size |  | Distortion (proportional area) |  |  |  |  |  | Calibrated estimate of dot diameter (um) |  |  |  | Proportional LPMM (line pairs per millimeter) |  |  |  |  |  |
| --- | --- | --- | --- | --- | --- | --- | --- | --- | --- | --- | --- | --- | --- | --- | --- | --- | --- | --- |
|  | Pixels | um | C | C- UL | C- UR | C- LL | C- LR | Mean Residual | C | Δ actual | Mean of corners | Δ actual | C | C- UL | C- UR | C- LL | C- LR | Mean Residual |
| **OCi** | 3024 X 3024 | 6661 X 6661 | 1.0 | 0.911 | 0.909 | 0.937 | 0.963 | 0.930 | 249 | -1 | 239 | -11 | 1.0 | 0.536 | 0.536 | 0.445 | 0.445 | 0.490 |
| **XPi** | 3024 X 3024 | 6403 X 6403 | 1.0 | 1.008 | 1.006 | 1.014 | 1.005 | 1.008 | 242 | -8 | 243 | -7 | 1.0 | 0.900 | 0.800 | 0.950 | 0.950 | 0.900 |
| **XP**s | 2976 X 2976 | 6485 X 6485 | 1.0 | 1.026 | 1.032 | 1.021 | 1.031 | 1.027 | 240 | -10 | 243 | -7 | 1.0 | 0.947 | 0.842 | 0.895 | 0.895 | 0.895 |
| **XT** | 2048 X 2048 | 6360 X 6360 | 1.0 | 1.012 | 0.996 | 1.022 | 1.002 | 1.008 | 251 | 1 | 252 | 2 | 1.0 | 0.889 | 0.889 | 0.889 | 0.889 | 0.889 |

The calibrated estimate of dot diameter determined from images was closest to the true value for center patch in the XT and the OCi, differing only by 1 um. For the OCi, the corner patches differed by 11 um, the worst performance of any configuration tested here, and the largest center-to-corner difference. The XT performed the best with the estimated diameter at the center patch only 1 um greater than actual, and with the corner estimates only 1 um larger than center. The XPi had a similar proportional difference between center and corners, but with a larger initial underestimate (8 um) of diameter in the center patch.

The maximum resolution in line pairs per millimeter (lpmm) in the center of an image was 101 for the OCi, 90 for the XT, 80 for the XPi, and 57 for the XPs. Using these values as the reference values, the comparison to the corners of the images (Table 1) showed that the XT and two XP configurations retained the largest proportion of optical resolution (89-90%) and therefore had the lowest spherical aberration. The OCi configuration retained less than 50% of its center-image resolution.

## Discussion

The images produced by the XyloPhone, especially the XPi are, for most metrics and to the human eye, as good as XT images (Figure 4, Figure 5). Distortion, while undesirable, can be removed using post-processing algorithms if the distortion for the device is mapped (Zhang 1999; Hartley and Kang 2005) or can be removed using machine learning approaches (e.g. Li et al. 2019). Systematic errors in estimating feature size (e.g. dot diameter) can be readily calibrated as well. Unlike geometric distortion in an image, spherical aberration cannot as easily be ameliorated. Based on the results here, the OCi configuration showed the largest change in resolution across an image and therefore would be a poor candidate for acquiring research-grade or archival macroscopic images, and may need additional post-processing for robust and stable training of image classification models for CVWID applications, especially if models were trained to use patches of the parent image. Any patch incorporating a corner of such an image will be incorporating tissue imaged at less than half the spatial resolution, so otherwise macroscopically observable features might be obscured.

Of the phone-based configurations, the XPi had the highest initial and residual resolution in the corners, suggesting that it would be best-suited for acquiring research-grade or archival macroscopic images with comparatively consistent parameters across the image. Given the similarity to XT image metrics and the existence of several published CVWID XT models (Ravindran et al. 2018, Ravindran et al. 2019, Ravindran et al. 2020, Ravindran and Wiedenhoeft 2020, Arévalo et al. submitted), the XyloPhone is clearly attractive hardware for a possible CVWID system. It is important to revisit the fact that the Samsung phone used in this study was a relatively old Samsung Note 5, whereas the iPhone was a much newer model. Because the Ollo Clip was evaluated on the same phone as the XyloPhone, comparisons between these systems are valid, and the XyloPhone greatly out-performed the Ollo Clip (Table 1), even when the Ollo Clip has the benefit of the XyloPhone illumination array.

I chose not to evaluate the Xylorix lens system in this work. Available preview images posted on the Xylorix website are 800 × 800 pixels, and only the central 350-400 × 350-400 pixel patch shows sufficient image quality for evaluation, with spherical aberration obvious and extreme in the remaining three fourths of the image. This observation precluded the need to evaluate the optical properties of the Xylorix lens, as it is clearly inadequate for research grade imaging and seems to produce images from which only the center patch is likely to reliably show the underlying wood anatomical features.

Tests are underway to compare the cross-compatibility of XP and XT wood images for CVWID models for laboratory testing and field deployment. If XP images can be used to train models for deployment on the XT, and conversely, if existing or new XT models can be deployed on a smartphone with the XP hardware and suitable software, the field-deployability of CVWID technology can be greatly expanded, as not only is the per-unit cost reduced by a factor of 12, but there is no need for a laptop for deployment (another several hundred USD cost reduction). In such a future, the total kit needed for field use would be a sharp utility knife, one’s own smartphone, and the XyloPhone.

In addition to its potential for application in CVWID systems, the XyloPhone can be used to image a wide range of biological or manmade materials (Figure 6), just as with the XyloTron (Ravindran et al. 2020). In principle, any substance with useful or interesting macroscopic variation can be imaged, if it can withstand contact of the XyloPhone, or if the user has a steady hand to image without contact.

**Figure 6.**
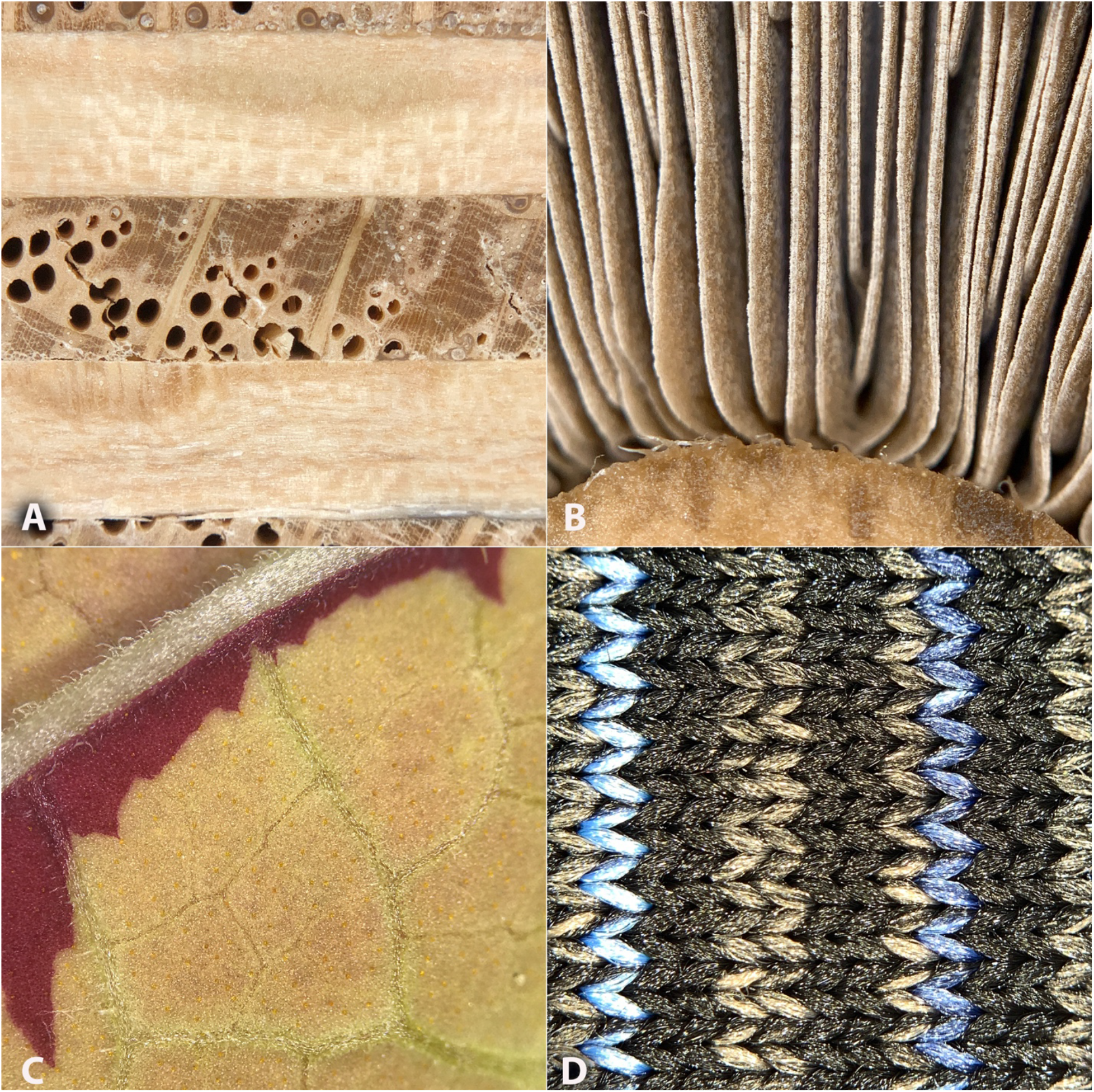
Free-hand visible light XyloPhone images of: A, red oak plywood. B, the stipe and gills of a white button mushroom. C, the abaxial surface of a variegated *Coleus* leaf. D, fabric from a sportswear shirt. A, B, and D are 6,742 um on a side. C, courtesy of Caitlin Gilly, using an iPhone 7 Plus, is 6,159 um on a side.

Open-source, affordable, high-performance imaging hardware like the XyloPhone can provide a consistent foundation from which one can build macroscopic reference data sets, but specimen selection and specimen preparation, at least for wood, become paramount, as they are likely to be the quality-limiting-factors in a dataset. Models developed from poor specimen sampling or inadequate hardware may represent cases of garbage-in fiction out (GIFO), a subset of the larger and well-known garbage-in garbage-out (GIGO) principle in computer science, and should give rise to healthy skepticism about the validity of any models built from such data sets. It is critical that as more research teams become involved in CVWID research, wood anatomists with key domain knowledge remain involved in the process. When specimens are poorly chosen, inadequately prepared, inappropriately imaged, and/or when the forensic questions are ill-conceived, the final result should be regarded warily (e.g. Olschofsky and Kohl 2020
). Hasty or poorly executed work has the potential to harm the advancement of the field by making astute readers skeptical of future CVWID research outputs and limiting its adoption. It is my hope that by providing an even-lower-cost alternative to the XyloTron (which remains as the gold-standard for wood imaging, albeit at a higher price-point), the XyloPhone can ensure that future data sets collected across the world share cross-compatibility that gives rise to a future with a master data set that can be selected from and deployed globally.

## Supporting information

Assembly manual and bill of materials

Contains STL files for 3D printing

Gerber files for ordering PCBs

## Acknowledgments

My sincere thank you to Dr. Blaise Thompson for providing the gerber files for the VIS LED PCBs. Special thank you to Dr. Prabu Ravindran for a few waves of detailed comments that greatly improved the structure and quality of the manuscript. My gratitude to Caitlin Gilly for the image of the *Coleus* leaf, and to the encouragement of the CWAR team more broadly. The 2020 COVID-19 quarantine allowed me the opportunity to develop and refine this device.

## Supplementary Information

- 3D files (XyloPhone main unit, charger cap, surface plate, electronics lid, spacer ring, a generic phone mounting plate, and several phone-specific plates)
- Assembly manual with a bill of materials
- Gerber files for PCBs – the power input, output, and distribution boards are as in Ravindran et al. 2020, and the VIS LED boards are adapted from those in that publication

