## Supplementary material for "The XyloPhone: democratizing access to high-quality macroscopic imaging for wood and other substrates": Assembly manual and bill of materials

### Constructing a XyloPhone

Alex C. Wiedenhoeft
Caitlin Gilly

Center for Wood Anatomy Research, Forest Products Laboratory
US Forest Service, US Department of Agriculture

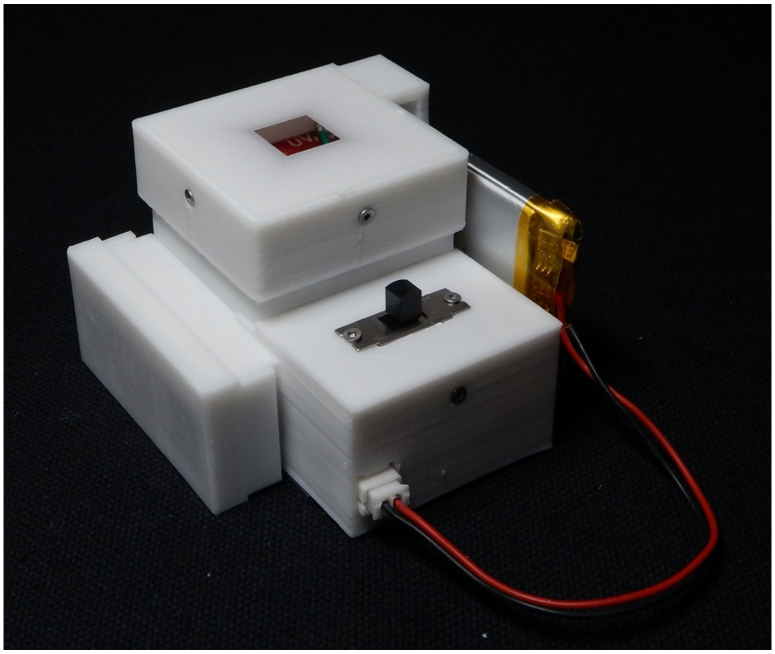


August 2020

Those portions of this manual recycled from the XyloTron assembly manual by Wiedenhoeft and Kleinschmidt (2020) are used here with permission of the authors.

#### TABLE OF CONTENTS

Part One: The Need for and Use of This Manual

Part Two: Bill of Materials

Part Three: Illustrated Parts List and Preparatory Work

Part Four: Step-by-Step XyloPhone Assembly and Calibration Instructions

#### PART ONE: THE NEED FOR AND USE OF THIS MANUAL

The XyloPhone you are about to build is the next generation of imaging hardware for the broader XyloTron platform, leveraging the electronics of the XyloScope, but in a compact new design to accommodate bring-your-own-device functionality. The XyloPhone's design reflects the intent that this device can be built by an enterprising end-user to image a wide range of substrates – it is an assemblage of individually simple off-the-shelf (itemized in the bill of materials in Part Two) and 3D printable components. With a phone-specific adapter it can be used on any smartphone.

In Part Three we present an illustrated parts list and a short description of preparatory work that needs to be done prior to assembly. In this section we also note with an * several parts for which it is prudent to acquire more than the minimum number, because, if a mistake is made and the part is ruined, having spares facilitates the efficient completion of your XyloPhone. Fortunately, those components most likely to be ruined in the learning process are among the most affordable.

The step-by-step instructions in Part Four are relatively detailed. Despite this, the manual does assume certain basic skills. Specifically, no instruction on the use of basic tools (screwdrivers, cutting tools) is provided, nor is there any tutorial in stripping wires or soldering electronic components. Such tutorials abound (e.g. learn.sparkfun.com). We further assume basic functionality in 3D printing and that, if a printed component is faulty or damaged, such components can be readily reprinted.

This manual is intended to provide detailed instructions to assemble the XyloPhone in its current incarnation. It also lays the groundwork for a novice to make special modifications themselves to adapt the XyloPhone for the widest possible breadth of uses. We look forward to a future where other workers develop and share designs superior to what we provide here, and we hope to benefit from those advancements.

Each part or component is given a number in the illustrated part list. In the assembly instructions, the list number of each part is presented in parentheses beside the part's name in the text so that should one experience any confusion, it can easily be remedied by consulting the list.

#### PART TWO: BILL OF MATERIALS

The bill of materials to build one XyloPhone. Minimum required quantity (Qty), XyloPhone part name, Vendor, Manufacturer (MFR), and manufacturer part number (MFR #) are listed. Please refer to the notes in Part Three for those parts for which acquiring more than the minimum possible number might be advisable. One will also need a smartphone, and to adapt a holder for that phone.

| Qty | XyloPhone part | Vendor | MFR | MFR # |
|---|---|---|---|---|
| 1 | LED controller | Digikey | Recom Power | RCD-24-0.30 |
| 1 | Voltage reference | Digikey | Analog Devices Inc. | AD680JTZ |
| 1 | Radial capacitor | Digikey | Murata Electronics | RDER71H104K0P1H03B |
| 4 | White LED | Digikey | Lite-On Inc. | LTPL-P00DWS57 |
| 4 | UV LED | Digikey | SunLED | XZVS54S-9A |
| 1 | SPDT switch | Digikey | NKK Switches | MS13AFG01 |
| 2 | Right angle JST Through-Hole 2-Pin Connector | Digikey | JST Sales America | S2B-PH-K-S(LF)(SN) |
| 1 | JST jumper two wire assembly 6" | Digikey | Sparkfun Industries | PRT-09914 |
| 4 | 1/2" No. 2 screw | McMasterCarr | McMasterCarr | 95893A560 |
| 7 | 1/4" No. 0 screw | McMasterCarr | McMasterCarr | 95893A505 |
| 1 | 1/4" No. 2 screw | McMasterCarr | McMasterCarr | 95893A550 |
| 1 | Belomo 10x Triplet hand lens | belomostore.com | Belomo | |
| 1 | Lithium ion battery | SparkFun | Lithium Ion Battery - 1Ah | PRT-13813 ROHS |
| 1 | Battery charger | SparkFun | SparkFun LiPo Charger Plus | PRT-15217 |
| 1 | Reversible USB A to C Cable - 0.3m | SparkFun | SparkFun | CAB-15426 ROHS |
| 1 | Power distribution PCB | | | |
| 1 | Power input PCB | | | |
| 1 | Power output PCB | | | |
| 4 | LED PCB | | | |

#### Part Three: Numbered and Illustrated Parts List and Preparatory Work

##### Numbered and Illustrated Parts List (number required within parentheses, * denotes prudence in acquiring more than the minimum number). Images are not to scale.

1. 3D printed XyloPhone housing

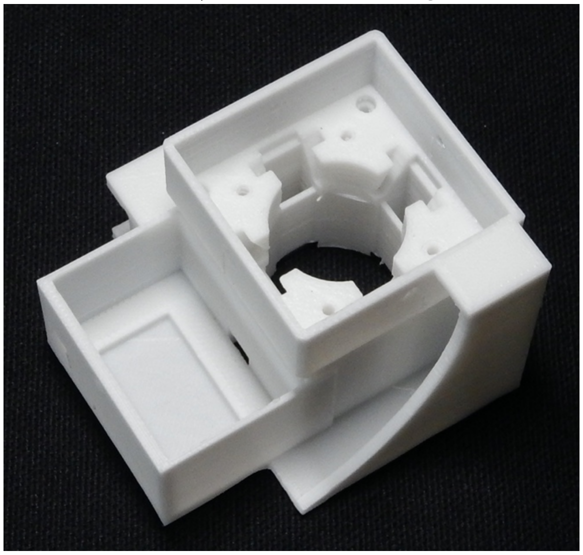

2. 3D printed charger cap

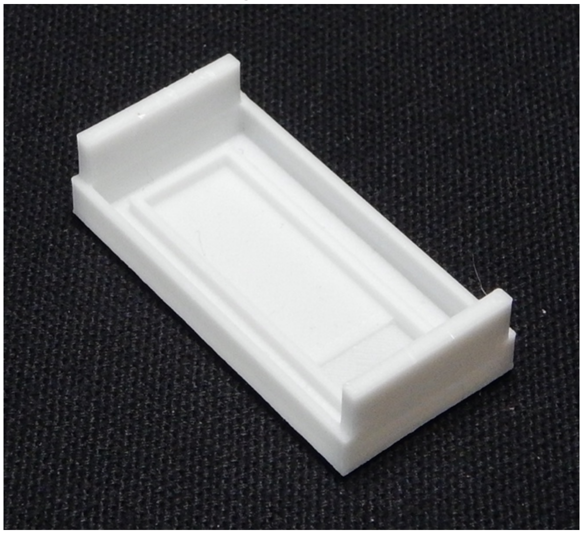

3. 3D printed electronics lid

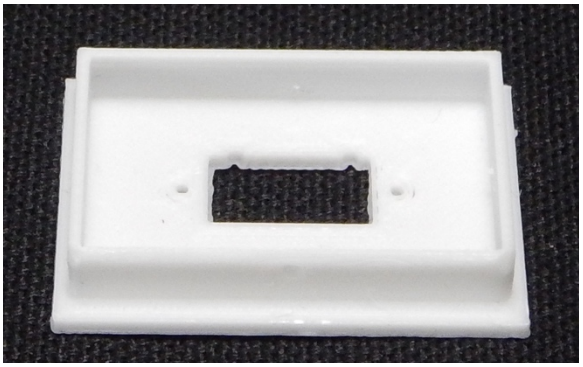

4. 3D printed spacer ring

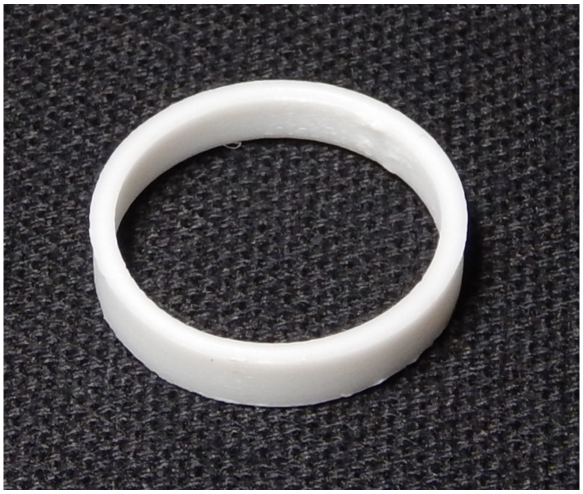

5. 3D printed surface cap

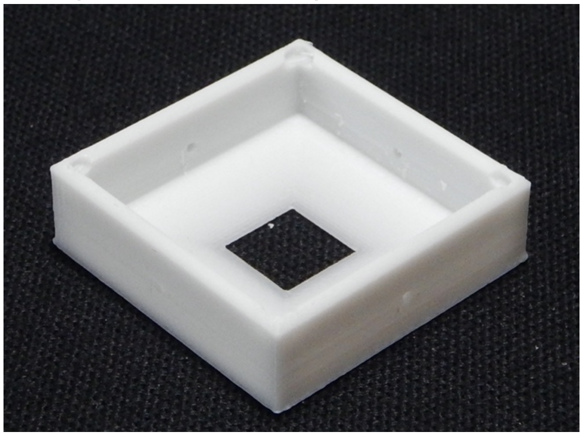

6. Battery

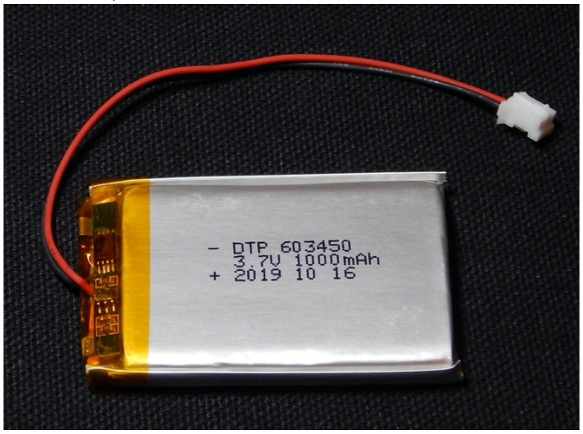


7. Battery charger

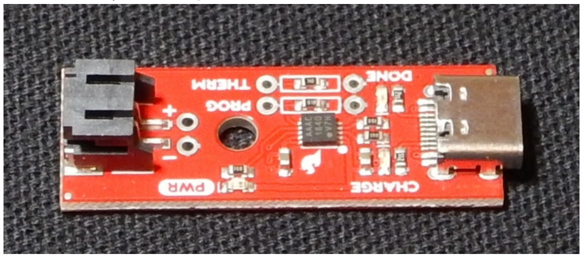


8. Charger cord

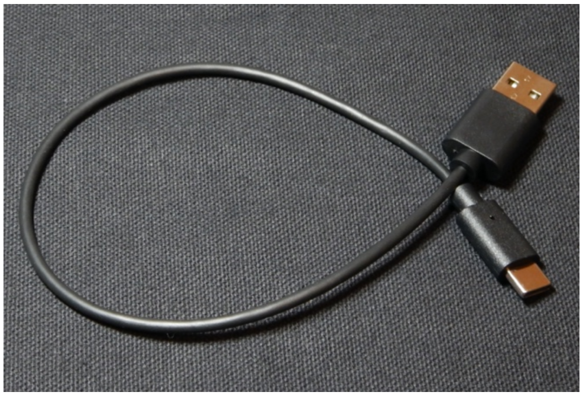

9. Belomo 10 X triplet loupe

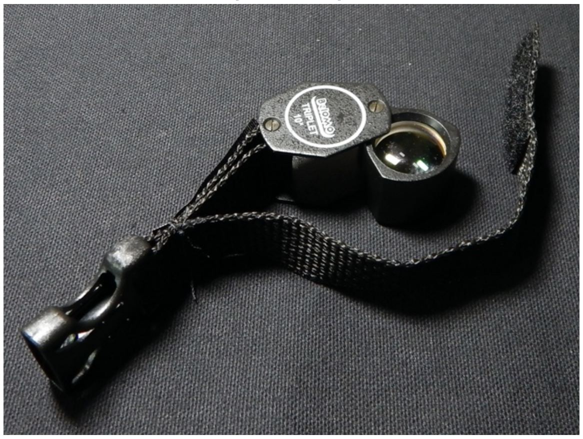

10. UV LED (4*)

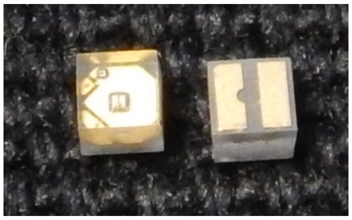

11. LED controller (1)

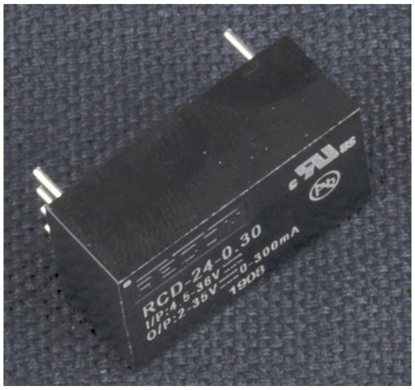


12. Radial capacitor (1)

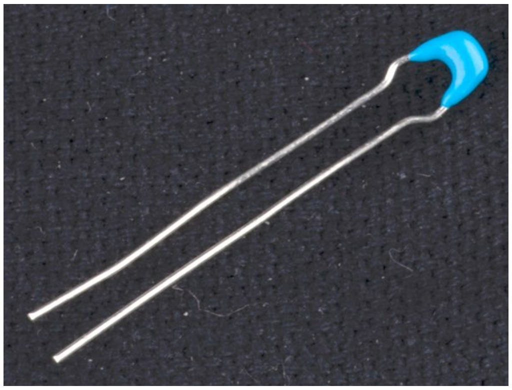

13. VIS LED (4*)

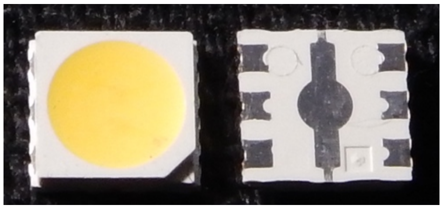

14. Switch (1)

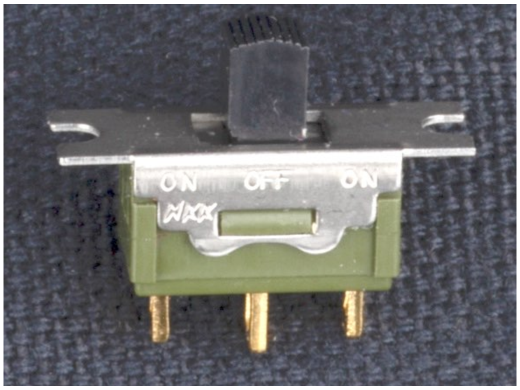


15. Right angle JST Through-Hole Connector (1*)

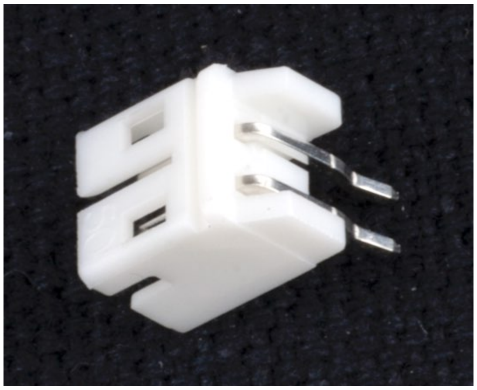

16. (a, b, c)
(a) 1/2” No. 2 screw
(b) 1/4” No. 2 screw
(c) 1/4” No. 0 screw

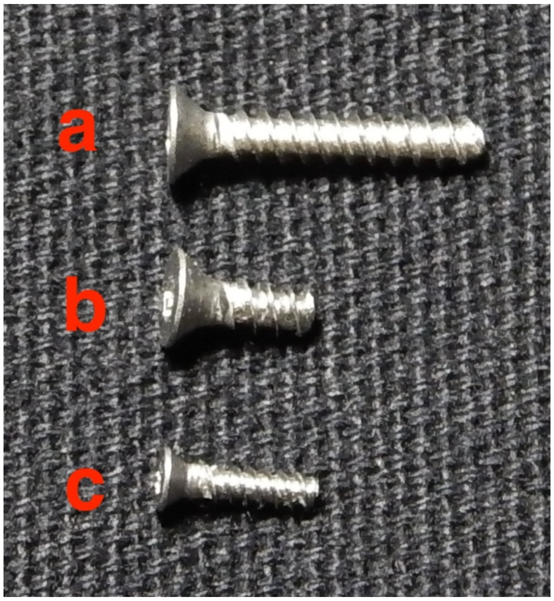


17.  Voltage reference (1)

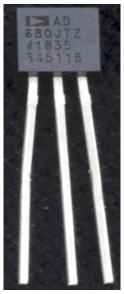


18. JST jumper two wire assembly 6" (1)

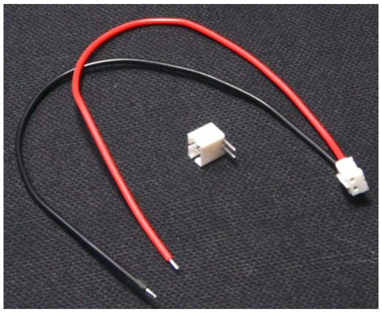

19. Power distribution PCB – top view only (1*)

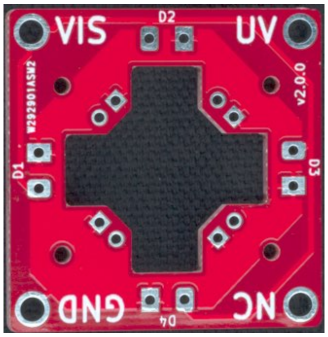


20. Power input PCB – top view, bottom view (1*)

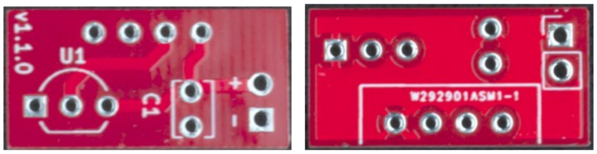


21. Power output PCB– top view, bottom view (1*)

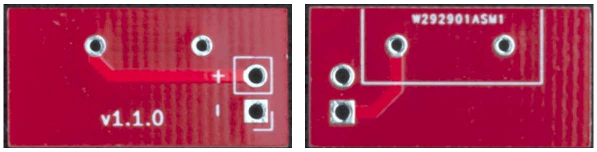

22. LED PCB (4*)

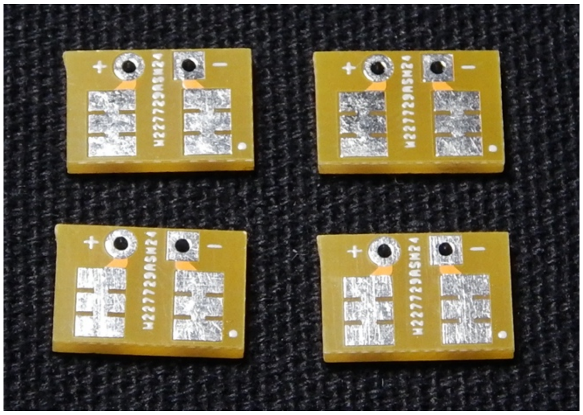


Note that for all PCBs, + through holes are round or have rounded corners, and – through holes are squares or have right-angle corners.

#### Preparatory Work

Print the 3D printed components. Clean them of any adhesive used in the printing process, and also do any trimming necessary - it is common that the plate face of the 3D printed component will have a lip that is slightly wider than the rest of the part (shown below via the end view of a freshly printed object printed in black polymer). This can be removed with a utility knife or a deburring tool.

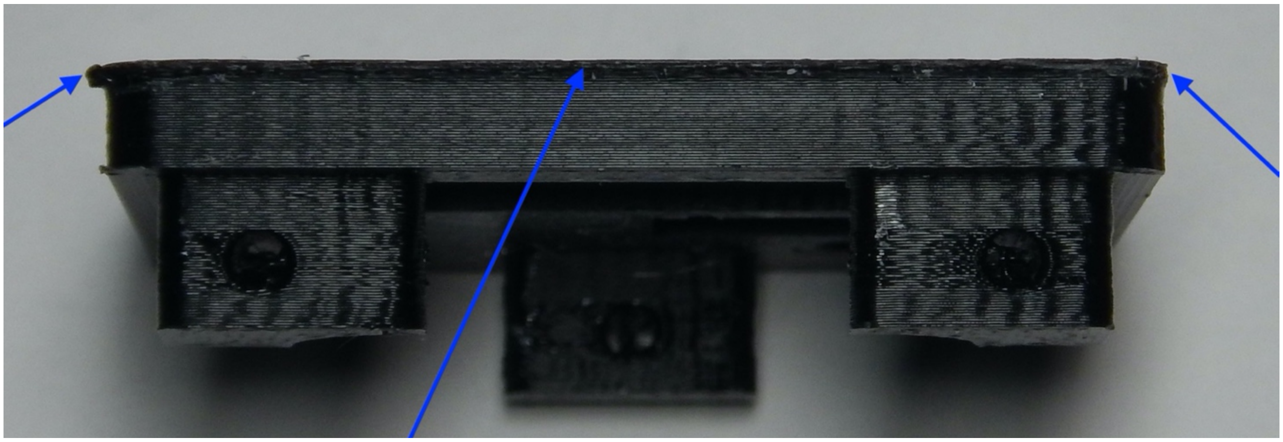

In the illustrations of this manual, there are only five 3D printed components, all of which were printed in white polymer to facilitate photographing the assembly process. It is recommended that in practice all components be printed in black to prevent any white-balance distortion.

Submit the Gerber files to order the printed circuit boards. We have printed the boards in red and white and in tan, but other color combinations are possible. Perhaps the most conservative selection would be black boards with dark grey printing so that the boards themselves cannot cause white-balance distortion.

Remove the lens from the Belomo 10 X triplet loupe (9). Open the loupe and find the side that has the internal nut holding the lens in place. Using a tool like an internal snap ring pliers, unscrew the nut and remove the lens.

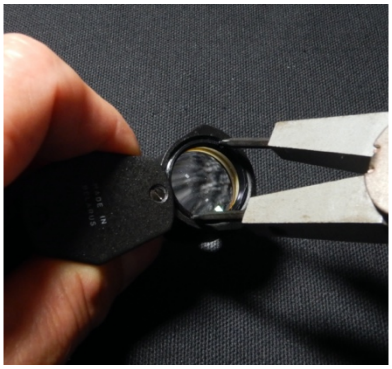

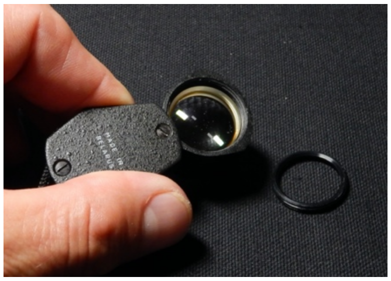

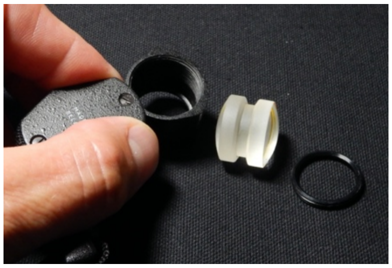

#### Part Four: Step-by-Step XyloPhone Assembly Instructions

##### Electronic components – soldering and wiring, phase I

*Power input assembly*

1. Clamp the LED controller (11) in a vise, pins-side up

2. Fit the power output PCB (21) onto the two pins on the LED controller (11). The power output PCB (21) has a partial rectangle printed to show where the controller seats – that printed side will be in contact with the LED controller (11), so there is only one way this board can be placed (note that the below left we are showing the face that will be in contact with the LED controller (11)). Below right we show the correct installation onto the two pins of the LED controller (11). Solder the two pins, then flush cut them.

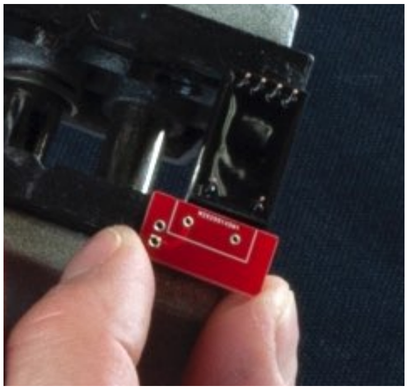
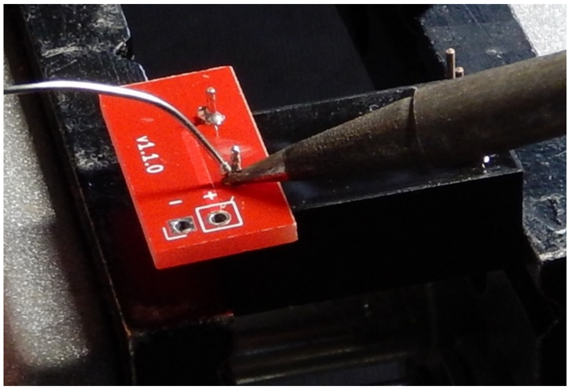


3. Fit the power input PCB (20) onto the four pins on the LED controller (11). As with the power output PCB (21), the power input PCB (20) has a partial rectangle printed to show where the LED controller (11) seats (note in the image below-left: we are showing the face that will be in contact with the LED controller (11)). Below-right: we show the correct installation onto the four pins of the LED controller (11) – note that the partial circle print is now facing up (lower right image). Solder all four pins, then flush cut them.

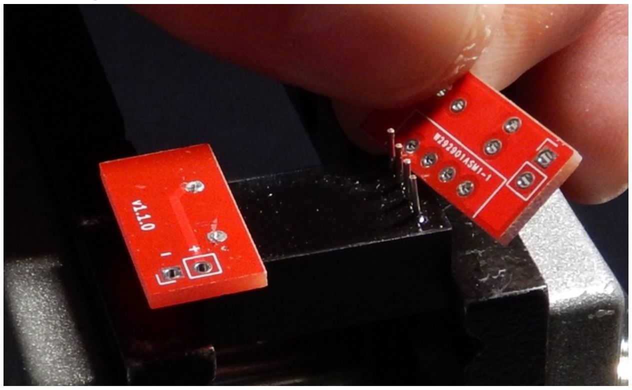

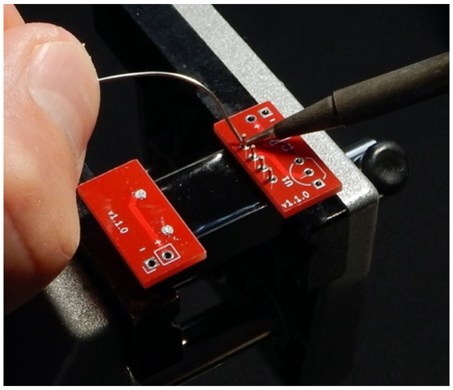

4. Insert the voltage reference (17) into the three holes in the power input PCB (20). There is a semicircular print with a flat edge associated with the three mounting holes on the board. Install the voltage reference (17) so that its flat face is facing the flat side of the shape outline. Gently and carefully bend the leads so that the bottom of the voltage reference (17) can be mounted within 2-3mm of the top surface of the power input PCB (20). Solder the three leads separately and flush cut them. If the voltage reference (17) is mounted too high, the assembly will not fit in in later steps.

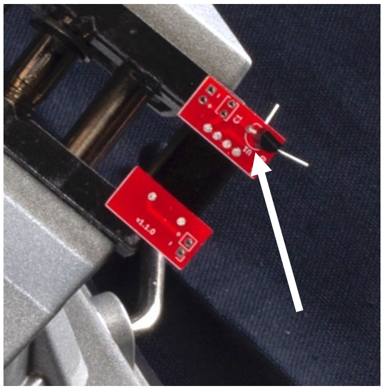

5. Insert the radial capacitor (12) into the two holes in the small printed rectangle (labeled C1) so that its height is less than that of the voltage reference (17). There is no special orientation for the radial capacitor (12). Solder and flush cut.

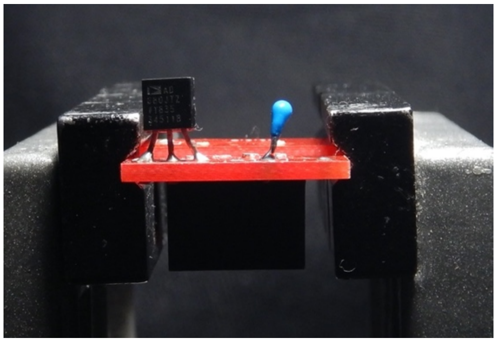

6. The last component of the power input PCB (20) is the right angle JST through-hole connector (15). This fitting is inserted from the LED controller (11) side of the power input PCB (20) so that its pins emerge by the printed – and + near the radial capacitor (12). Solder these two pins, taking care to make mechanically strong connections with the board without angling the pins or melting through to the plastic housing. Flush cut the pins.

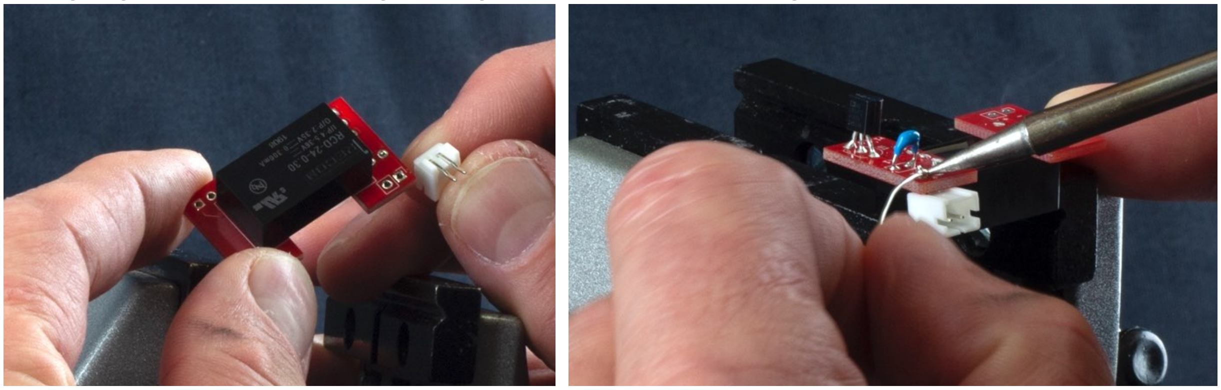

VIS LED-LED PCB assemblies

7. Pinch the VIS LED (13) onto the LED PCB (22) with flat tweezers, and then use a small binder clip to holder the tweezers in place. The compliance of the tweezers ensures that excessive force is not applied to the VIS LED but permits fine spatial adjustment of the VIS LED (13) position on the LED PCB (22). (The VIS LED (13) is best placed about 0.5mm from the outer edge of the LED PCB (22) and the small cut-out on the corner of the VIS LED (13) must face the outer edge of the – pad (left image, below)). This assembly is then clamped in the vise. Heating the soldering pads of the LED PCB (22) without touching the VIS LED (13) and then allowing capillary flow to make the connection is usually successful. It is important to use a minimal amount of solder so that the VIS LED-LED PCB assemblies will slide smoothly into the 3D printed XyloPhone housing (1). Repeat this process three more times, for a total of four completed VIS LED-LED PCB assemblies.

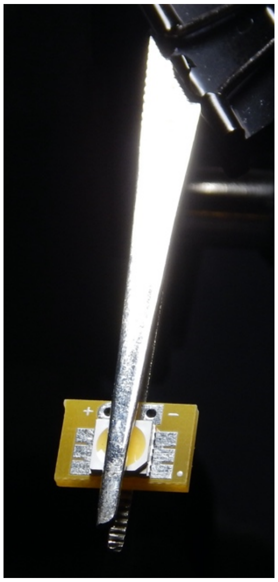

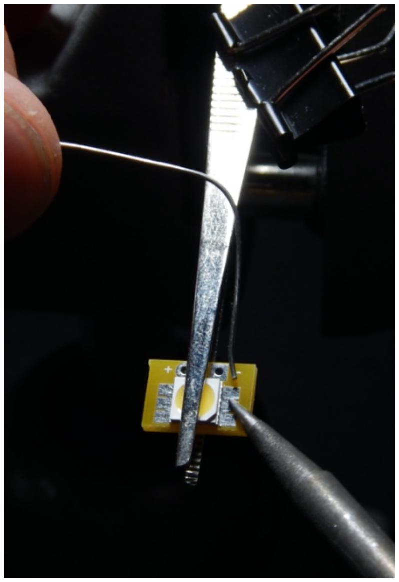

8. Using an LED testing setting on a multimeter, ensure that each VIS LED-LED PCB assembly lights <u>both</u> internal LEDs in the VIS LED (13). This is best done at the +/- through holes on the LED PCB (22), to ensure the entire assembly is electrically correct.

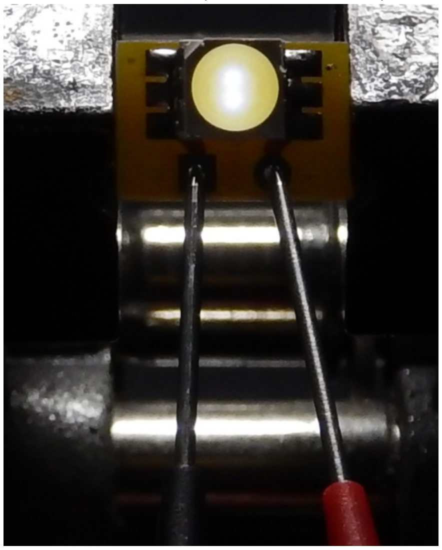

9. Insert the twisted stranded wire into each through hole on the VIS LED PCB (22). Solder the wires in the through holes. Flush cut the back side of the boards, then trim the excess wire to no less than 7mm.

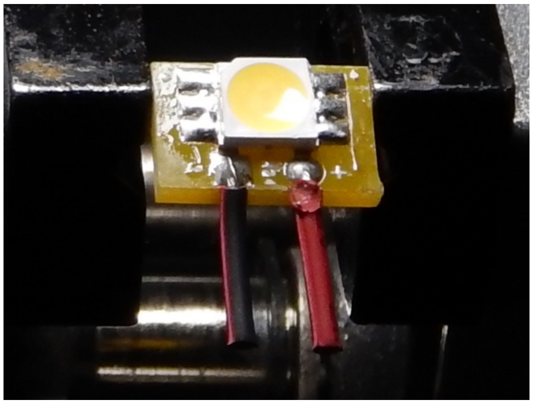

10. Twist the wire as you strip the insulation from the ca. 7mm of wire, ensuring that it is tightly twisted. Lightly tin this now-bare wire ensuring that that you do not increase the outer diameter of the braid- it must be able to pass through the through-holes in the power distribution PCB (19).

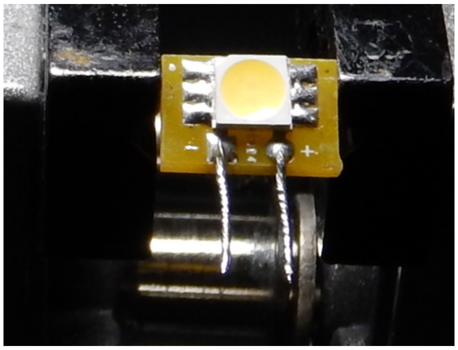

11. Given the difficulty of mounting the electronics into the 3D printed XyloPhone housing (1) in a later steps, it would be prudent to use a multimeter to test each VIS LED (as in step 8), using the electrodes on the tinned wires.  These connections will become inaccessible in later steps.

Power distribution PCB-UV LED assembly

12. Clamp the power distribution PCB (19) in the vise, with the side with the printed "UV" "VIS" "NC" and "GND" facing up.

    Note: You will be soldering the UV LEDs (10) to the pairs of though-holes on the diagonally-cut peninsulas adjacent to center opening in the power distribution PCB (19).

13. If you are right-handed then fill the right (and only the right) of the two through holes at each peninsula with solder.  There should be a slightly (but only slightly) raised meniscus of cooled solder in each through hole when you are done. If you are left handed, you would fill the left through-hole.

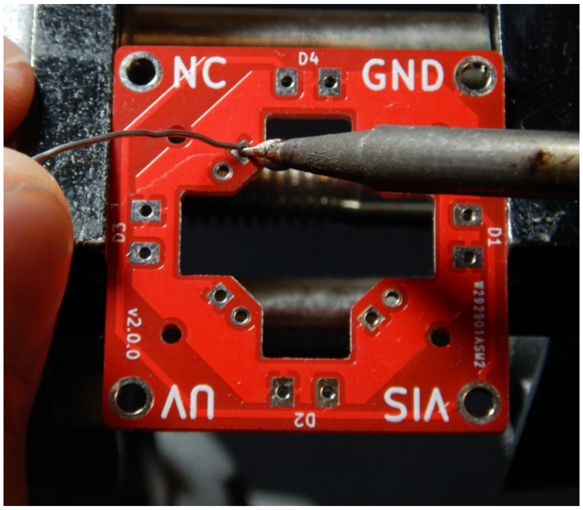


**THE NEXT TWO STEPS ARE THE MOST DIFFICULT OPERATIONS IN THE ASSEMBLY OF THE XYLOPHONE**

14. Orient the UV LED (10) so that its positive pad corresponds to the + through hole and so that you can press it flush to the face of the through hole.  Get a tiny bead of melted solder on the tip of the soldering iron and use that that to melt the solder in the through hole.  Lower the UV LED (10) in place, gently press it onto both through holes of the power distribution PCB (19), and then remove the soldering iron.  When the solder cools, one of the two pads of the UV LED (10) will be soldered.

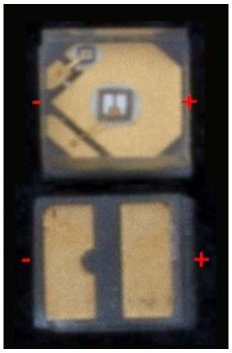

15. Place the solder at an angle into the through hole adjacent to the through-hole you just soldered to the UV LED (10), and WITHOUT TOUCHING THE LED WITH THE SOLDERING IRON, solder the other pad of the UV LED (10). Repeat these steps for the other three UV LEDs (10). The power distribution PCB-UV LED assembly is complete for now.

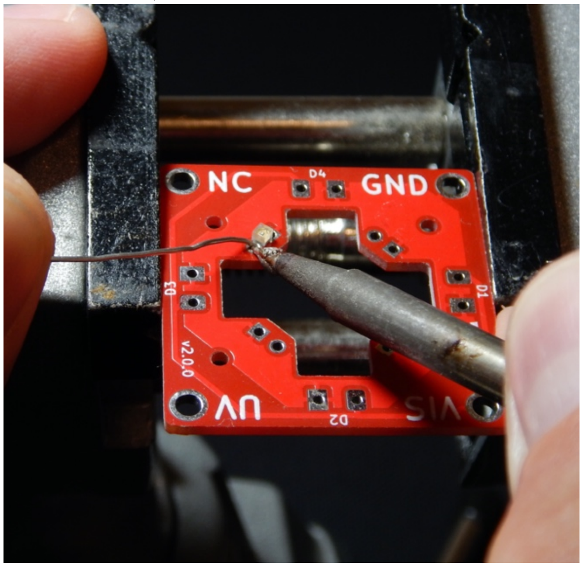


16. Before proceeding, it is a good idea to confirm that your soldering work is sound and electrical connections are made correctly. This is most easily done by using the power input assembly completed in Step 6 to power the power distribution PCB-UV LED assembly. Plug the JST male connector of the Battery (6) into the female JST connector (15) on the power input board (20). There is but one way to accomplish this, so it is not pictured.

17. Using the JST jumper two wire assembly (18), push the pins through the through holes on the power output PCB (21) ensuring that the red wire is connected to the + through hole and the black wire is connected to the - through hole.

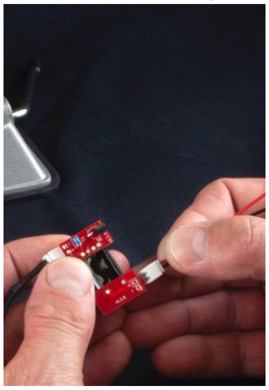

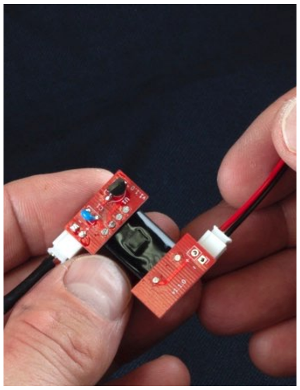

18. Ensuring that the pins of the JST jumper two wire assembly (18) are in contact with the through holes of the power output PCB (21), touch the black (-) wire to the GND corner's large through hole, and the red (+) wire to the UV corner's large through hole on the power distribution PCB-UV LED assembly. If all electrical connections are correct, all four UV LEDs (10) will light up. Remove the JST jumper two wire assembly (18) if all is well. If one or more UV LEDs (10) fail to light, you can work to re-solder the unlit UV LEDs (10). Iterate on this process until all four UV LEDs (10) light.

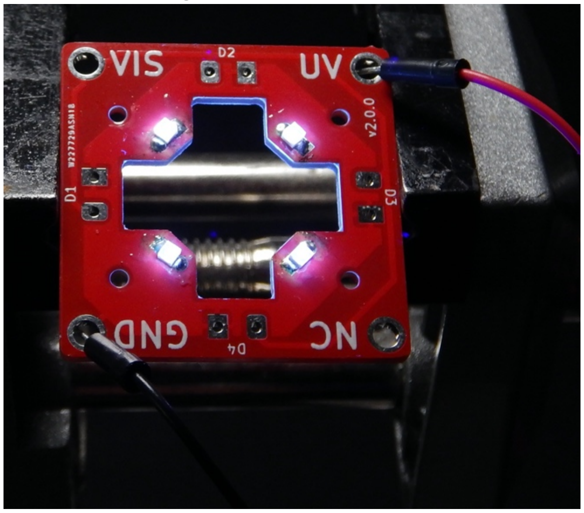


**MECHANICAL ASSEMBLY – PHASE I**

#### LED holder-power distribution PCB assembly

19. Now that the UV LEDs (10) are soldered onto the power distribution PCB-UV LED assembly **it is important that as you handle the assembly you do not damage the UV LEDs** (10). They are well-suited to thousands of hours of data collection, but are not resilient in the face of mechanical damage. Insert each of the four VIS LED-LED PCB assemblies into the bottom face of the power distribution PCB (19) such that the VIS LED (13) is facing the center cut-out in the power distribution PCB (19). This will ensure that the – and + terminals on each of the boards match.

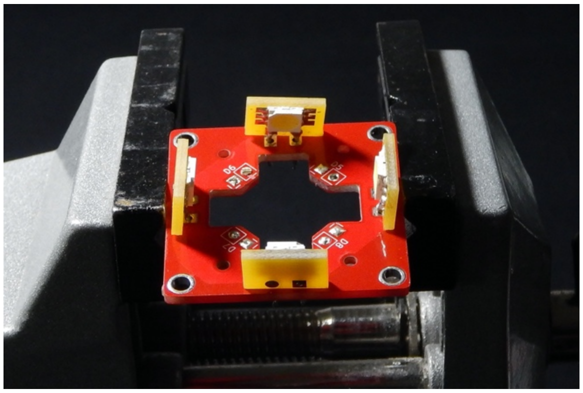

**Read this entire step prior to starting, as board orientation with regard to the 3D printed XyloPhone housing (1) is critical**

20. Using great care not to damage the UV LEDs (10), tip the 3D printed XyloPhone housing (1) sideways, and push the VIS LED – LED PCB -power distribution PCB-UV LEDs assembly into the large rectangular manifold at the top of the 3D printed XyloPhone housing (1). The VIS LED-LED PCBs will enter the manifold first, and each will slide into an angled slot to orient them correctly. Each board will need to be manipulated separately into its slot, and this process is made easier by the horizontal orientation of the 3D printed XyloPhone housing (1) and can be facilitated by the judicious use of fine-tipped forceps. The ~7mm of wire soldered to each VIS LED-LED PCB should provide ample play to situate each assembly without it slipping from the power distribution PCB. The rotation of the power distribution PCB with regard to the 3D printed XyloPhone housing (1) is critical. We have found it easiest to orient the board as depicted below, but please note this orientation (vs. 180 degrees, which the only other functional option) necessitates a longer UV and VIS power supply wire path in later steps. In my mind, we have preferred this longer path length and a power distribution PCB orientation that favors the printed text of UV and VIS on the corners of the power distribution PCB in an "upright" position, but that is an entirely biased view based on my preferred (and arbitrary) "up" orientation for the 3D printed XyloPhone housing (1). (A wiring parsimonist would clearly prefer a frame of reference inverted relative to my preference, and given those assumptions, they are completely correct.) Once the VIS LED -LED PCBs are fully seated in their slots, carefully but forcefully press the power distribution PCB into the manifold until it is flush across its entire face. *It is all too easy to damage the UV LEDs in this phase*. Ensure that the holes in the power distribution PCB nearest the UV LEDs are aligned with the holes in the 3D printed XyloPhone housing (1). These are necessary for routing power in subsequent steps. If these holes are not aligned, the power distribution PCB and VIS LED-LED PCB assemblies will need to be carefully extracted, rotated 90 degrees, and re-installed.

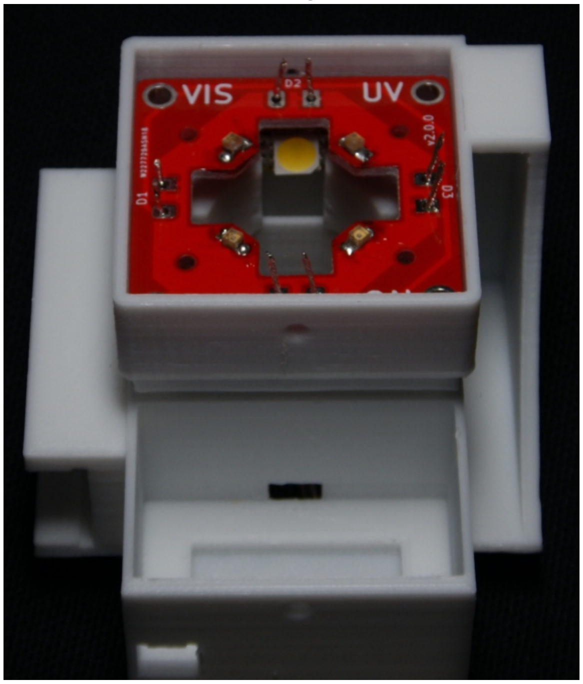

**ELECTRONIC COMPONENTS – SOLDERING AND WIRING, PHASE II**

#### LED holder-power distribution PCB assembly

21. The ~7mm wires soldered to each VIS LED board are now protruding from the top face of the power distribution PCB. Taking great care not to touch the soldering iron to the 3D printed manifold, solder each wire in the through hole and flush cut.

22. As with Steps 16-18, it is prudent (and satisfying) to confirm that the electrical connections are sound. Using the same assembly in Steps 16-18, and ensuring that the pins of the JST jumper two wire assembly (18) are in contact with the through holes of the power output PCB (21), touch the black (-) wire to the GND corner's large through hole, and the red (+) wire to the VIS corner's large through hole on the power distribution PCB-UV LED assembly. If all electrical connections are correct, all four VIS LEDs (13) will light up. Remove the JST jumper two wire assembly (18) if all is well. If one or more VIS LEDs (13) do not light, check the connections at the through-holes to the power distribution PCB (19).

**WIRING THE POWER DISTRIBUTION PCB**

23. Through the small holes near each UV LED (10), feed a narrow (e.g. 28 ga. stranded insulated) wire through the holes adjacent to the GND, UV, and VIS large corner through-holes. The wires must be long enough (e.g. ~16 cm) to reach the corner through holes on the top of the 3D printed XyloPhone housing (1), and must be long enough to route through the channels in the base, and into the electronics compartment on the front of the 3D printed XyloPhone housing (1).

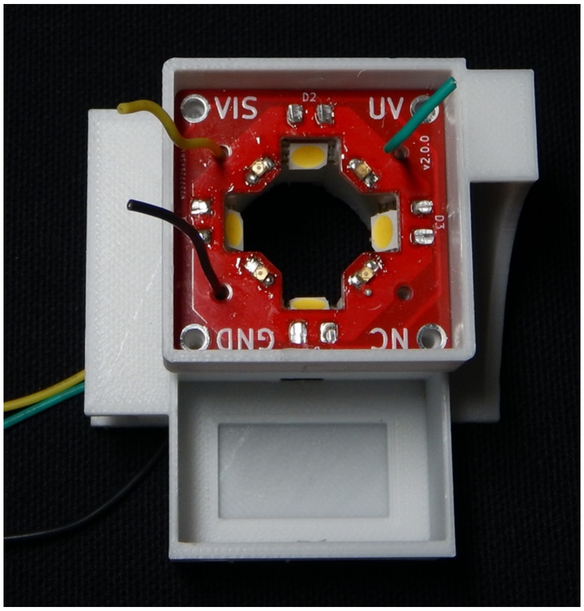

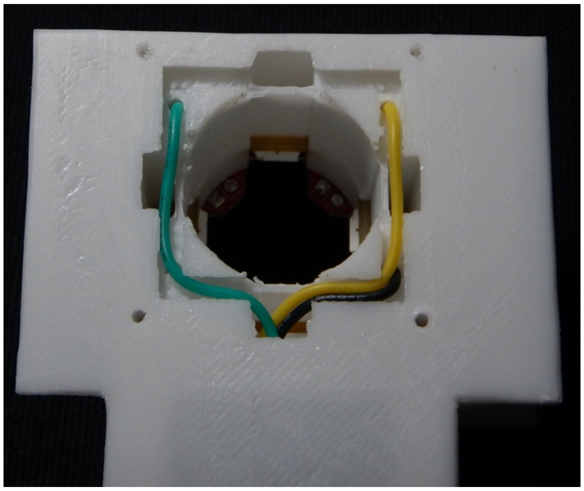

24. Strip insulation from the end of each wire, twist the end, and form a small coil or ball that can be tucked into the large corner through hole in the power distribution PCB (19). Taking great care not to melt the manifold, solder each wire to the corner through holes.

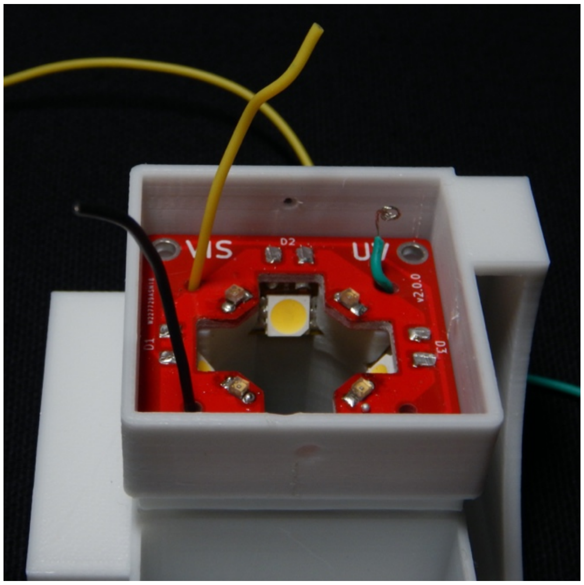

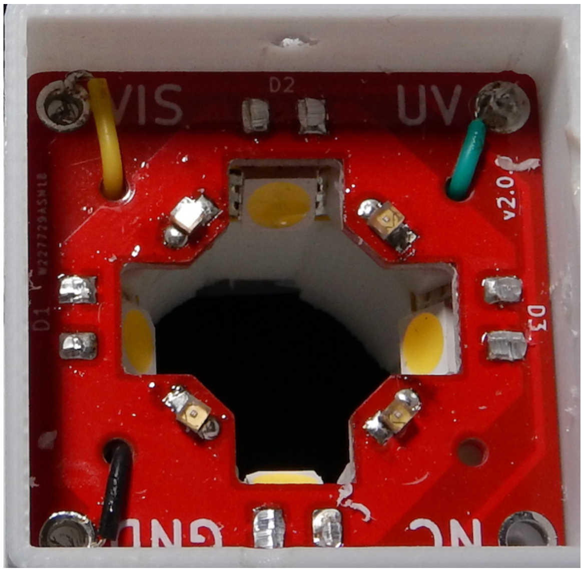


25. Flip over the 3D printed XyloPhone housing (1) and route the wires through the channels in the base, and out the small opening into the electronics compartment. Securing the wires with an even layer of clear tape that does not obscure the central cylindrical opening is convenient. When the wires are routed correctly they will look something like what is shown below (arrow indicating the clear tape).

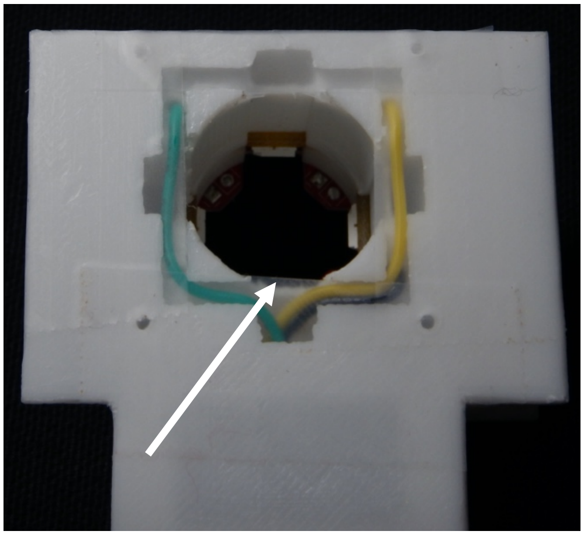

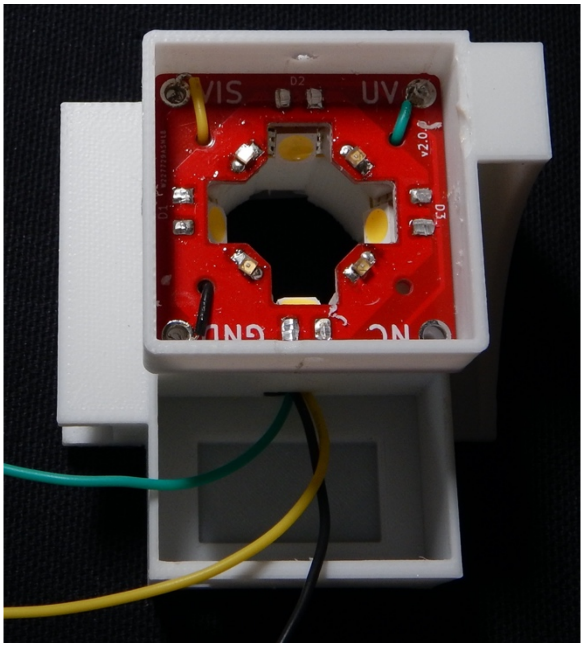

26. Install the power input assembly in the electronics compartment. Pull aside the wires and angle the assembly so that the JST connector (15) can protrude through the printed hole in the outer wall of the electronics compartment. The LED controller (11) will snap into the printed recess in the base of the compartment. It will be a snug fit, and lightly sanding off or trimming the right angle of the rear edge of the LED controller (11) can facilitate this.

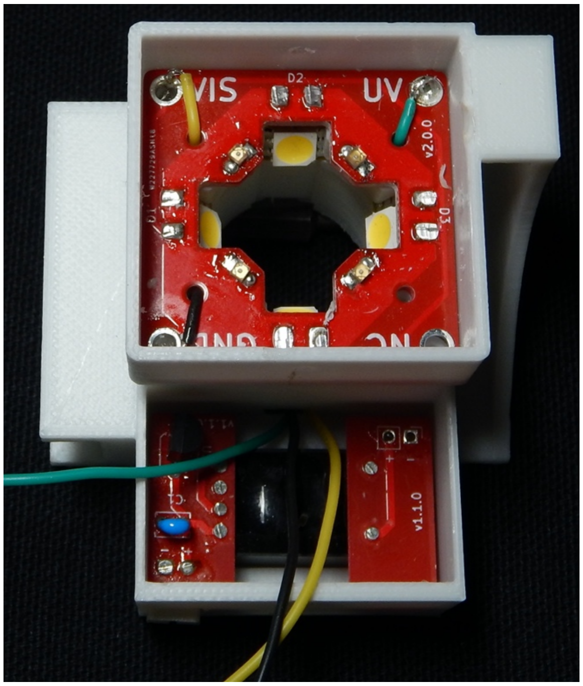


27. Wire the GND wire (black in the images here) to the – through hole on the power output PCB (21). Trim the black to wire to a suitable length, strip, twist and solder.

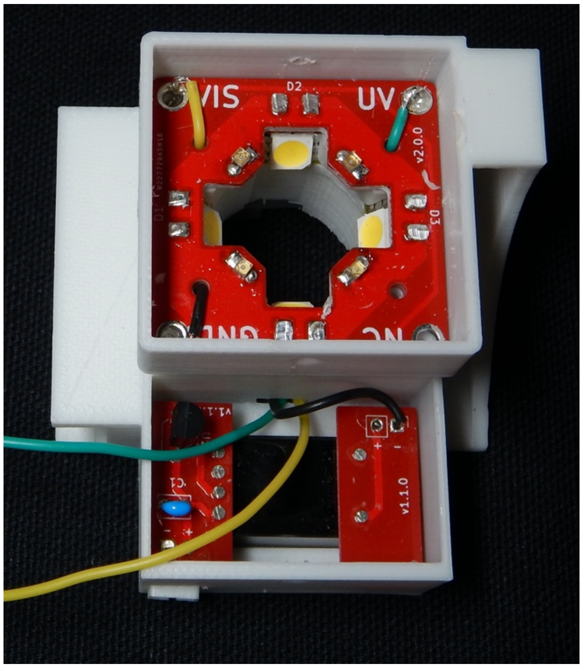


28. Cut a suitable length of wire (e.g. ~4cm) to be soldered to the + through hole on the power output PCB (21) – we used a red wire. This wire will connect to the center terminal on the switch (14). Strip, twist and solder to the + through hole.

29. Fit the switch (14) into the printed slot in the 3D printed electronics lid (3), then slide one piece of shrink tubing up each of the three wires – do not let them come too close to the switch (15) or the heat of soldering will cause premature shrinkage. Trim the wires as short as will permit you to wire them to the switch (14) and still manipulate the 3D printed electronics lid (3). Strip the end of the wire connected to the + terminal of the power output PCB (21)(red in the images here), twist, pull it through the center terminal on the switch (14), mechanically twist it to the terminal, then solder in place. Repeat this wire stripping and soldering for the remaining two wires (one yellow "VIS" and one green "UV") to the remaining two lateral terminals. Allow the wires and posts to cool. Slide the shrink tubing down over the cooled, soldered terminals. Carefully shrink the tubing with a heat gun – the heat gun can easily melt 3D printed electronics lid (3) or the 3D printed XyloPhone housing (1) so some care is needed.

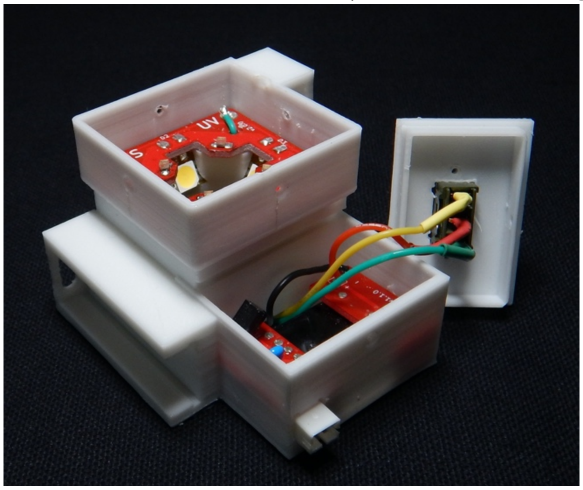


30. Snap the 3D printed electronics lid (3) into place, taking care that the shallower lip is seated in the slot in the 3D printed XyloPhone housing (1). It requires some force to make this fit. Secure the 3D printed electronics lid (3) with a ¼" No. 0 screw (16c), and use two more to secure the switch.

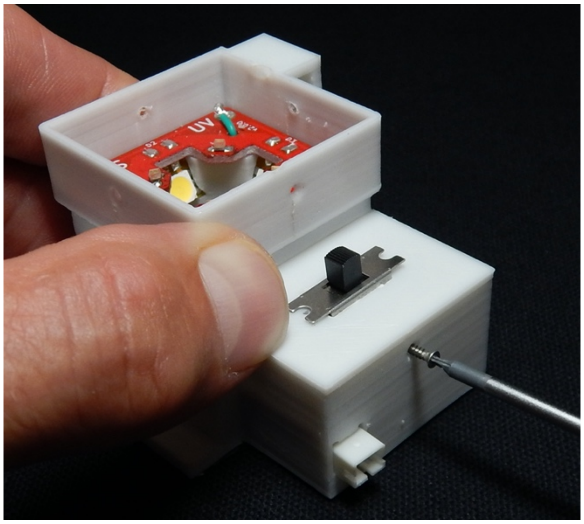


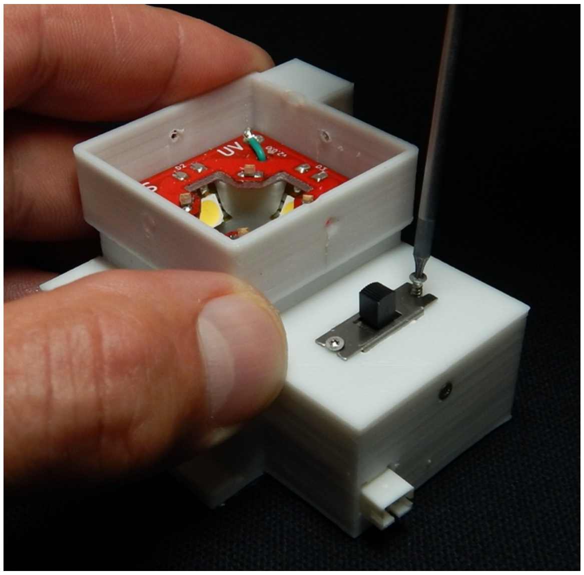

31. Install the 3D printed surface cap (5) into the manifold in 3D printed XyloPhone housing (1). If the power distribution PCB (19) was installed perfectly flush with the internal supports, the 3D printed surface cap (5) should snap in similarly flush. The fit is quite snug, and in reality there is probably little need for it, but using four ¼" No. 0 screws (16c), secure the 3D printed surface cap (5).

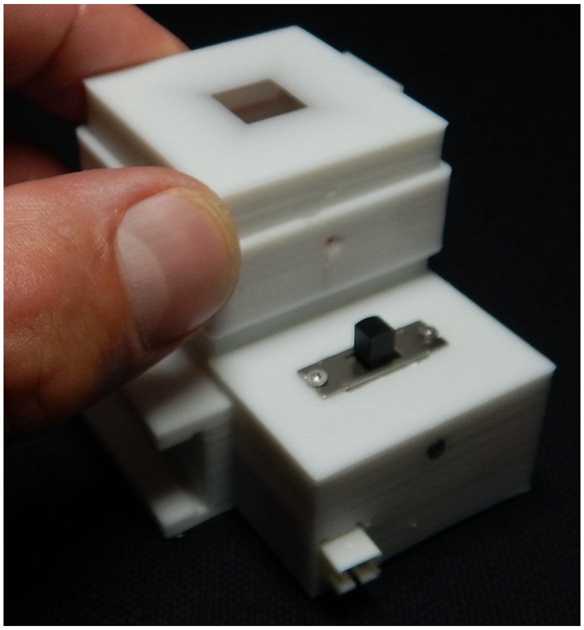

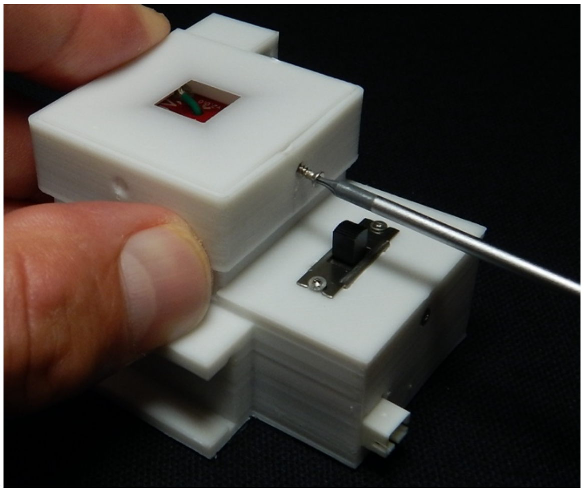

32. Slide the battery charger (7) into the charger slot in the 3D printed XyloPhone housing (1). The oval USB-C end of the battery charger (7) slides in first to mate with the oval printed hole. When the charger is fully seated, secure it with a ¼" No, 2 screw (16b).

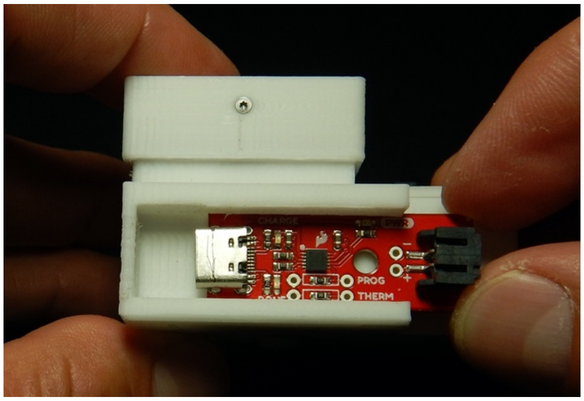


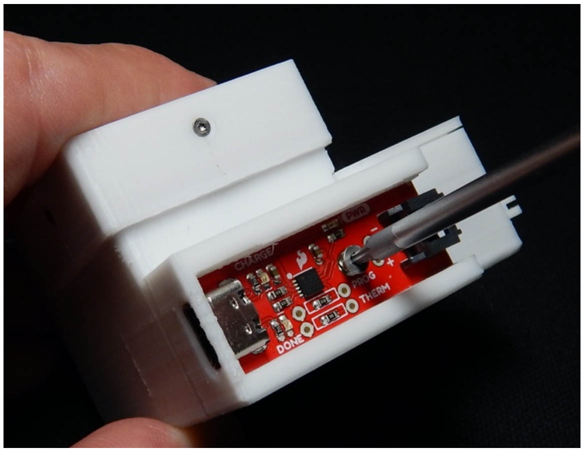


33. Snap the charger cover over the charger. Note that one end is shallower to allow for the JST connector, and at the other end the internal rim of the cover allows for a snug fit at the USB-C end.

34. Fit the battery (6) with one or more rubber bands of thickness sufficient to ensure that it will not inadvertently slip from the battery slot on the side of 3D printed XyloPhone housing (1). Slide the battery into the slot. Plug the male JST connect from the battery into the power input JST on the electronics compartment for use, or remove the charger cap and plug it into the charger if the battery needs to be charged. The charger cord (8) plugs in at the USB-C end, and can be charged with any typical 5V cell phone charging block or USD port on a computer.

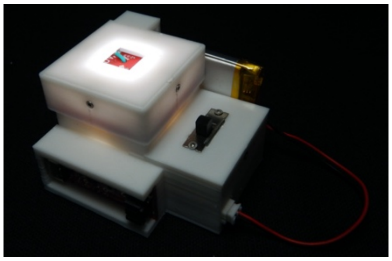
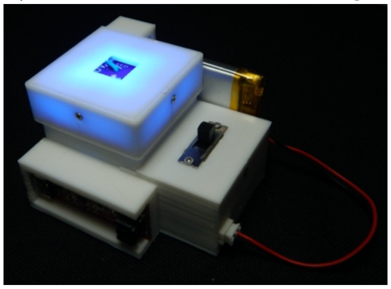
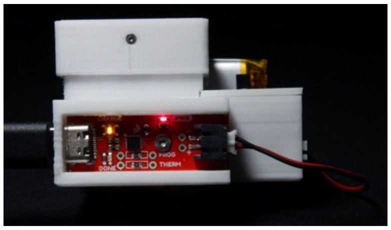

35. Ensuring that the lens from the Belomo 10X triplet loupe (9) is free from fingerprint and dust on both faces, place it into the central cylindrical opening in the 3D printed XyloPhone housing (1). Press it into place until it reaches the 3D printed stops.

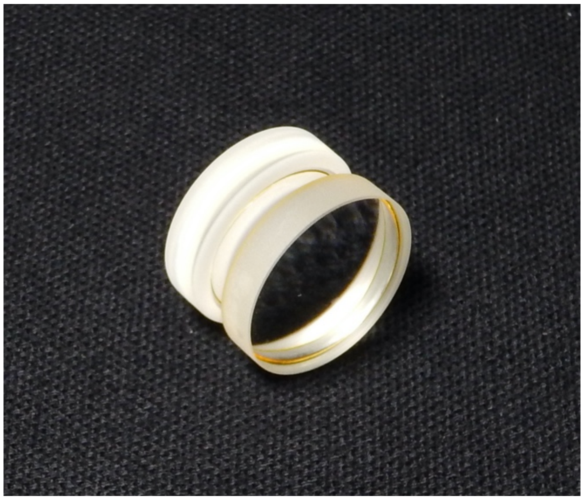
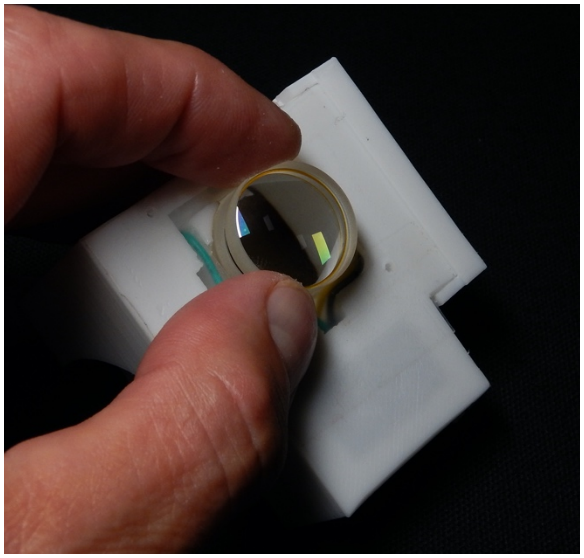

36. Place the 3D printed spacer ring (4) into the central cylindrical opening in the 3D printed XyloPhone housing (1). Press it into place firmly.

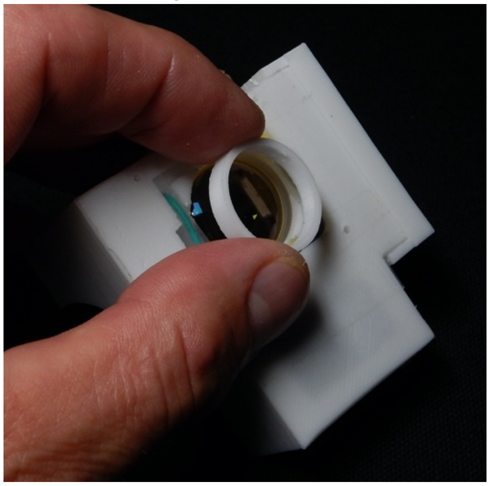 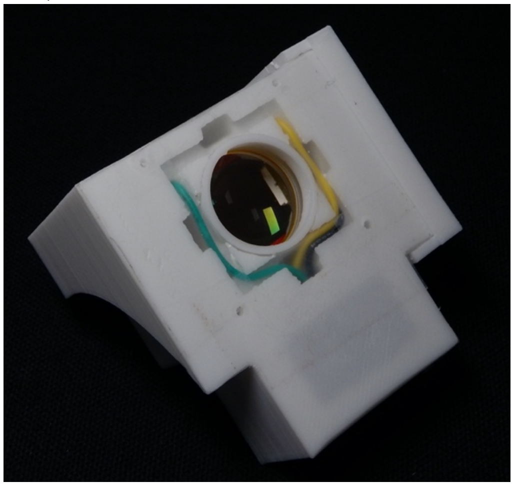

37. This completes the XyloPhone imaging and lighting unit.  What remains is to attach the XyloPhone to a customized base or mount for your phone.  We have provided the basic 3D file for the XyloPhone-to-smartphone interface; the center point of the four mounting screws holes is the center of the light path in the XyloPhone.  It is incumbent on the user to develop a phone-specific interface.  Below we show two such interfaces.  One slides over the phone with no case, is held in place by friction, and is good for lab work or extended use of the XyloPhone for data collection.  The other is a “paddle” design that fits into the smarthphone case, but must be held in place by the user.  This is convenient for intermittent and field use where it is helpful to keep the phone protected in its case.

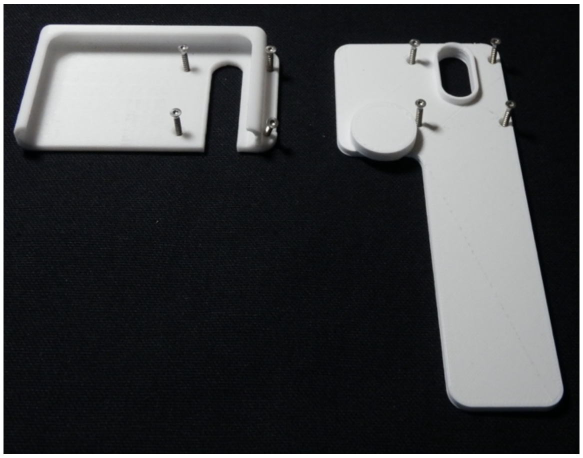 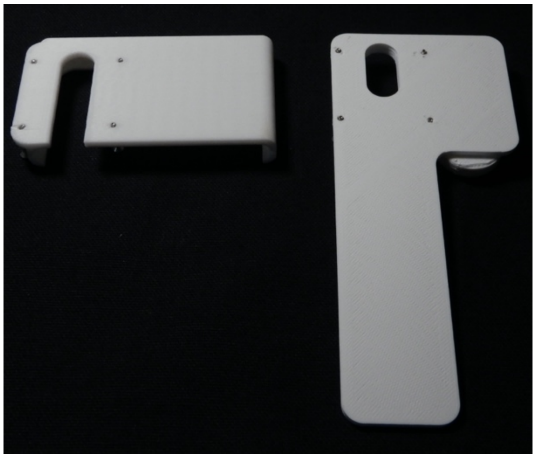

38. Mounting the XyloPhone to either base is the same process; using four ½" No. 2 screws (16a), screw the XyloPhone onto the base, being careful not to overtighten the screw and strip the plastic in the XyloPhone. Depending on the orientation of the XyloPhone and the design of the phone base, it can be necessary to adjust the design of the 3D printed charger cap (2) so that the lower edge of the cap does not interfere with the base.

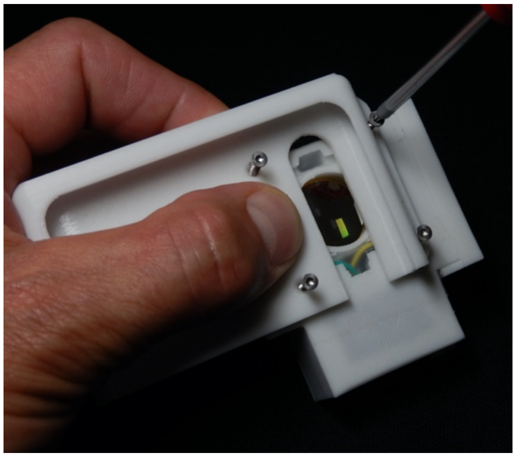

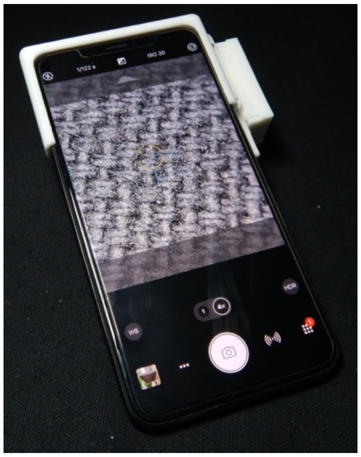

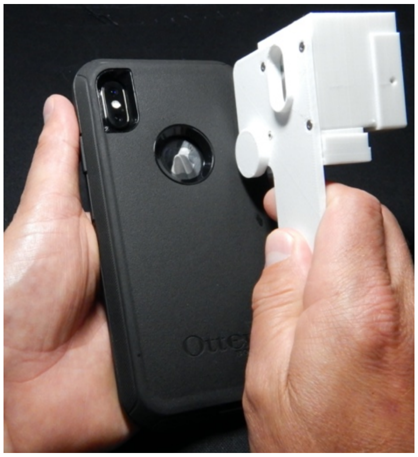

**Note that depending on the lens-switching parameters of your specific phone, the default camera app native to your operating system may not suffice to operate the XyloPhone; you may need to purchase or otherwise acquire a third-party camera application that allows you to control the camera(s) as needed.**

You have constructed a new XyloPhone – your skills are complete.
